# Modeling the motion of disease-associated KIF1A heterodimers

**DOI:** 10.1101/2023.06.22.546060

**Authors:** Tomoki Kita, Kazuo Sasaki, Shinsuke Niwa

## Abstract

KIF1A is a member of the kinesin-3 family motor protein that transports synaptic vesicle precursors in axons. Mutations in the *Kif1a* gene cause neuronal diseases. Most patients are heterozygous and have both mutated and intact KIF1A alleles, suggesting that heterodimers composed of wild-type KIF1A and mutant KIF1A are likely involved in pathogenesis. In this study, we propose mathematical models to describe the motility of KIF1A heterodimers composed of wild-type KIF1A and mutant KIF1A. Our models precisely describe run length, run time, and velocity of KIF1A heterodimers using a few parameters obtained from two homodimers. The independent head model is a simple hand-over-hand model in which stepping and detachment rates from a microtubule of each head are identical to those in the respective homodimers. Although the velocities of heterodimers expected from the independent head model were in good agreement with the experimental results, this model underestimated the run lengths and run times of some heterodimeric motors. To address this discrepancy, we propose the coordinated head model, in which we hypothesize a tethered head, in addition to a microtubule-binding head, contributes to microtubule binding in a vulnerable one-head-bound state. The run lengths and run times of the KIF1A heterodimers predicted by the coordinated head model matched well with experimental results, suggesting a possibility that the tethered head affects the microtubule binding of KIF1A. Our models provide insights into how each head contributes to the processive movement of KIF1A and can be used to estimate motile parameters of KIF1A heterodimers.

**SIGNIFICANCE:** KIF1A is responsible for transporting synaptic vesicle precursors in axons. KIF1A mutations are associated with neurodegenerative diseases. Most of these mutations are de novo and autosomal dominant, suggesting that half of the motors in patients are heterodimers composed of wild-type and mutant KIF1A. However, reliable theoretical models to explain the behavior of heterodimeric motors are lacking. In this study, we obtained exact analytical solutions to describe run length, run time, and velocity of heterodimeric motors which move in a hand-over-hand fashion. Our models provide valuable tools for quantitatively understanding the impact of heterodimerization with mutant KIF1A and the cooperative behavior of KIF1A dimers.

## INTRODUCTION

Axonal transport is fundamental for neuronal function and driven by motor proteins that move processively and directionally along microtubules (1). Kinesin superfamily proteins (KIFs) are molecular motors for anterograde transport (2). Among KIFs, KIF1A, a kinesin-3 family member, transports a synaptic vesicle precursor to synapses along the axon (3–5). KIF1A is monomeric in solution; however, KIF1A forms a dimer on cargo vesicles for efficient anterograde axonal transport (6, 7).

In humans, the motor domain of KIF1A is a hot spot that is associated with congenital neuropathies (8–10). The neuropathies caused by KIF1A mutations are called KIF1A-associated neurological disorder (KAND) (9). KAND mutations are mostly de novo and autosomal dominant. While a few mutations are gain of function mutations, most KAND mutations are loss of function mutations (9–16). We and others have suggested that loss of function KIF1A mutations significantly impair the motility of heterodimeric motors and axonal transport (16, 17). While the dominant-negative effect of KAND mutations has been uncovered, the specific inhibitory mechanism resulting from heterodimerization with KAND mutant KIF1A remains poorly understood.

Prior works have established models that can describe the behavior of kinesin dimers (18). Kaseda *et al*. demonstrated that heterodimers composed of non-identical kinesin-1 exhibit alternating fast and slow 8 nm steps, probably corresponding to displacement by the wild-type and mutant heads, respectively (19). The mean velocity of heterodimeric kinesin-1 was accurately reproduced using the velocities of two types of motors (19). However, analytical solutions for other important parameters of heterodimeric motors, such as the run length and run time, have not yet been developed. For analysis of homodimeric motors, single exponential fitting is useful for obtaining the mean run length and run time, as these parameters are theoretically exponentially distributed (20). To assess whether this conventional method used to analyze homodimers is applicable to heterodimers, mathematical verification is necessary.

The purpose of this work is to propose simple models that can describe the stepping and dissociation motions observed for heterodimers composed of non-identical KIF1A. The models proposed in this study describe the run length, run time, and velocity of heterodimeric KIF1A using a few parameters obtained from homodimers. We obtained exact analytical solutions; therefore, our models can be readily used by experimenters to analyze their data. Our formulation allows us to quantitatively understand how dimerization of wild-type KIF1A with disease-associated mutants affects the motility of the motor. The other purpose of this study is to investigate the coordination between the two heads of KIF1A. Each head of heterodimeric kinesin-1 steps alternately and independently along the microtubule (19). On the other hand, a study on KIF3AC, a mammalian neuron-specific kinesin-2 and a heterodimer of KIF3A and KIF3C motors, has demonstrated that KIF3A can accelerate the steps of KIF3C (21). Although KIF1A is known for being the fastest and most processive motor among the three neuronal transport kinesin families (22–24), the coordination between the two heads has remained elusive. Our study showed that the heterodimeric KIF1A moves at the expected speed as each head steps alternately at its intrinsic rate. Interestingly, with regards to the processivity of KIF1A, our models suggest that not only a microtubule-bound head but also a tethered head significantly contributes.

## MATERIALS AND METHODS

### Model for homodimeric motors

To analyze homodimer data, we construct a theoretical model to describe the stepping and dissociation motion of KIF1A homodimers. Recent stopped-flow fluorescence spectroscopy and single-molecule assays have uncovered an ATP-dependent stepping cycle of KIF1A dimer at saturated concentrations of ATP (25) (Fig. 1). In state 0, one motor domain is weakly associated with the microtubule and the motor can detach from it; the detachment terminates the processive run (transition 0→ unbinds). It is often the case that the tethered head binds to its next binding site (transition 0→1) before the microtubule-bound head detaches. After releasing ADP from its front head in state 1, KIF1A tightly binds to the microtubule (transition 1→2). The rear head in state 2 unbinds from the microtubule by releasing P_i_ from this head (transition 2→3). ATP then binds to the bound head in state 3 (transition 3→4) and undergoes hydrolysis, triggering full neck linker docking, which positions the tethered head forward and puts the motor in the weakly bound state again (transition 4→0). Symbols *k*’s indicate rate constants corresponding to these transitions as shown in Fig. 1. Some reverse transitions (ADP-on and ATP-off) are not indicated in the cycle diagram because these transitions are expected to occur at much slower rates compared to their forward transitions under conditions of high ATP and no ADP concentrations (26, 27). Under no load, a backward step rarely occurs compared to a forward step (28); therefore, this reaction is not included in the model. In the main text of this article, we use the model for highly processive motors obtained using certain approximations. The solutions of the full model are provided in Section S1 of the Supporting Materials and Methods. Since *k*_*i*_ is much faster than *k*_0d_ for highly processive motors, the time taken to complete one cycle can be approximated as follows:

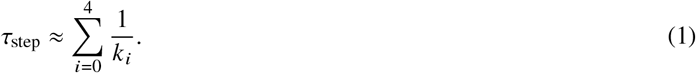

We will express the run length and run time distribution using *k*_0_, *k*_0d_ and *τ*_step_ (see Section S1 of the Supporting Materials and Methods and Fig. S1A). The run length *l* is defined as the distance traveled by the motor until its detachment. Let *a* be the motor step size (8 nm), and the run length distribution is given as

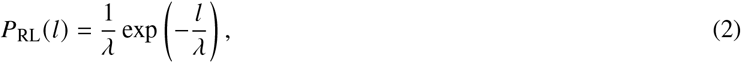

where *λ* defined by

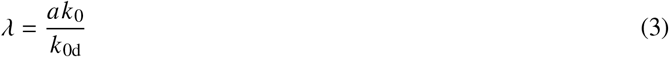

represents the mean run length. The expressions (2) and (3) are consistent with those used in previous studies (25, 29). The run time *t* is defined as the period of time that the motor remains on the microtubule until its detachment and its distribution is expressed as

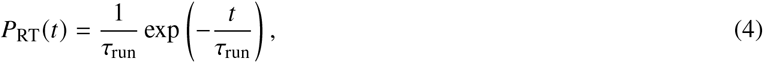

where *τ*_run_ defined by

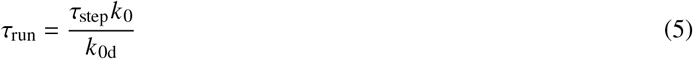

represents the mean run time.

**Figure 1:**
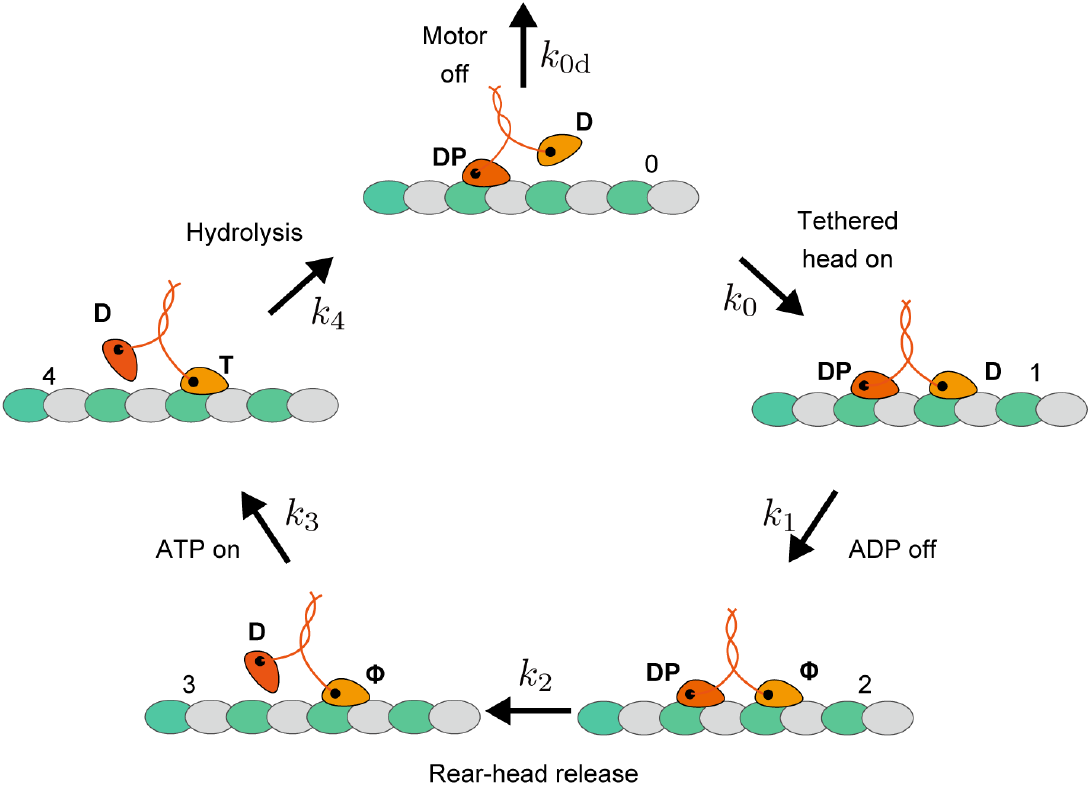
A model for homodimeric KIF1A. The diagram represents the stepping cycle for KIF1A homodimer under conditions with a saturated concentration of ATP (25). In the model, the cycle begins with one head bound to the microtubule weakly (state 0). From this vulnerable state, the bound head can detach from the microtubule and terminate the processive run (transition 0→ unbinds). Otherwise, the tethered head binds to its next binding site (transition 0→1). ADP is released to generate a tightly bound state (transition 1→2). The rear head releases P_i_ and detaches from the microtubule (transition 2→3). ATP binds to the microtubule-bound head (transition 3→4) and it is hydrolyzed to ADP-P_i_, which triggers full neck linker docking and positions the tethered head forward (transition 4→0). The motor is in the weak bound state again (state 0). *k* symbols indicate the rate constants. D, ADP; T, ATP; DP, ADP-P_i_; *ϕ*, apo.

### Model for heterodimeric motor

We construct a theoretical model to describe the motility of KIF1A heterodimers. We assume that KIF1A heterodimer moves in a hand-over-hand fashion. The heterodimer consists of two heads from different motors, motor A (orange head) and motor B (blue head), and a cycle consists of two forward steps, as shown in Fig. 2. The cycle in Fig. 2 begins when either head A or head B is weakly associated with the microtubule (state 0 or 5). In this state, the heterodimer can detach from the microtubule and terminates processive run (transition 0 → unbinds or →5 unbinds). Mostly, the tethered head binds to its next binding site (transition 0→1 or 5→ 6) before the bound head detaches. After releasing ADP from its front head, the motor tightly binds to the microtubule (transition 1→2 or 6→7). The rear head unbinds from the microtubule by releasing P_i_ (transition 2 → 3 or 7 → 8). ATP binds to the microtubule-bound head (transition 3 → 4 or 8 → 9) and is hydrolyzed, which induces full neck linker docking and positions the tethered head forward (transition 4 → 5 or 9 → 0). The motor is now in the vulnerable one-head-bound state (state 5 or 0) with the roles of the two heads exchanged compared to the starting state (state 0 or 5). From this state, the last half of the cycle starts, which proceeds similarly to the first half. Symbols *l*’s indicate rate constants corresponding to transitions in the cycle as shown in Fig. 2. As in the case of the model for homodimers, ADP-on, ATP-off, and backward step are not included in the cycle. In the main text of this article, we use the model for highly processive motors obtained using some approximations. The solutions of the full model are provided in Section S2 of the Supporting Materials and Methods. Since *l*_*i*_ is much faster than *l*_0d_ and *l*_5d_ for highly processive motors, the time for the heterodimer to complete two forward steps can be approximated as follows:

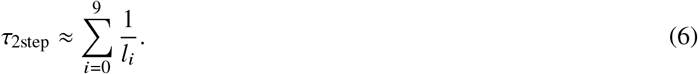

We will express the run length and run time distribution and the mean velocity of the heterodimeric motor using *l*_0_, *l*_5_, *l*_0d_, *l*_5d_ and *τ*_2step_ (see Section S2 of the Supporting Materials and Method and Fig. S1B).

The run length distribution of the heterodimer is given as

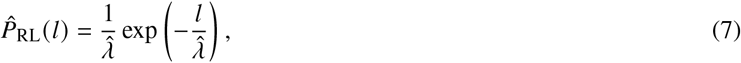

where

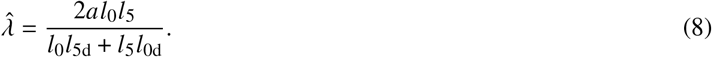

The run time distribution of the heterodimer is expressed as follows:

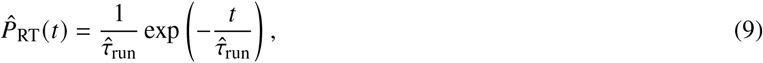

where

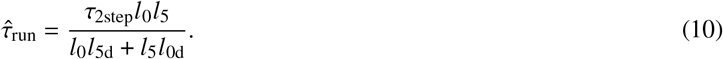

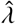 and 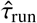 in Eqs. 8 and 10 represent the mean run length and run time of the heterodimer, respectively. Note that according to Eqs. 7 and 9, the run length and run time distributions of the heterodimer can be described by a single exponential function within the observable ranges of the run length and run time as explained in Section S2 of the Supporting Materials and Methods. The mean velocity of heterodimer is given by 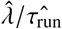 (30), which results in

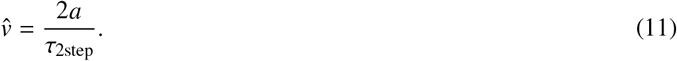

This expression is consistent with those used in previous studies (19, 31).

**Figure 2:**
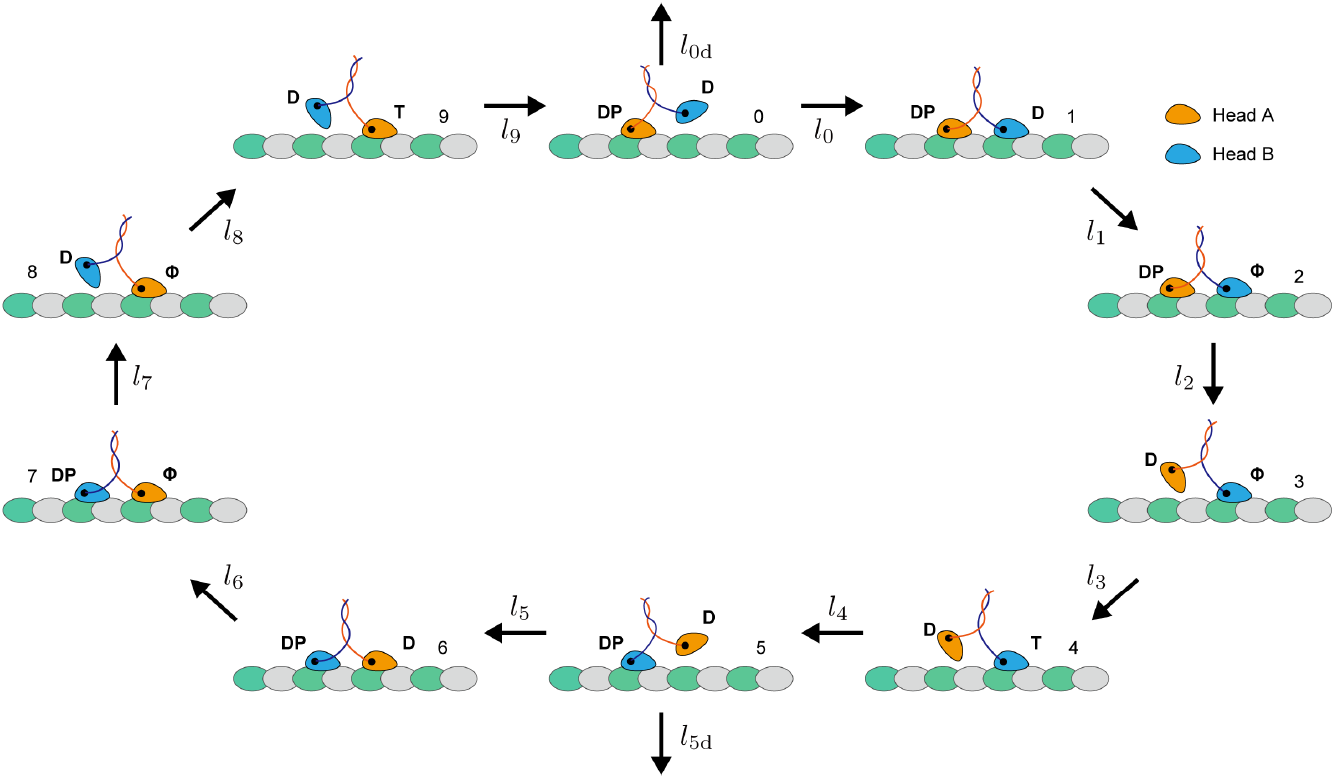
A model for heterodimeric KIF1A. The diagram represents the stepping cycle for KIF1A heterodimer under conditions with a saturated concentration of ATP. The heterodimer is composed of motor A (orange head) and motor B (blue head). In the model, the cycle begins when either head A or head B is bound to the microtubule weakly (state 0 or 5). From this vulnerable state, the bound head can detach from the microtubule and terminate the processive run (transition 0→ unbinds or 5→ unbinds). Otherwise, the tethered head binds to its next binding site (transition 0→1 or 5→ 6). ADP is released to generate a tightly bound state (transition 1→ 2 or 6→ 7). The rear head releases P_i_ and detaches from the microtubule (transition 2 → 3 or 7 → 8). ATP then binds to the microtubule-bound head (transition 3 → 4 or 8 → 9) and is hydrolyzed to ADP-P_i_, which triggers full neck linker docking and positions the tethered head forward (transition 4 → 5 or 9 → 0). The motor is in the vulnerable one-head-bound state (state 5 or 0) with an opposite head from the previous vulnerable state (state 0 or 5). *l* symbols indicate the rate constants. D, ADP; T, ATP; DP, ADP-P_i_; *ϕ*, apo.

### Protein Purification

We purified human KIF1A protein as described (16). Reagents were purchased from Nacarai tesque, unless described. Plasmids to express recombinant KIF1A are listed in Table S1. To directly study the motility parameters of the motors, the C-terminal regulatory and cargo binding domains were removed (Fig. S4 A and B). The neck coiled-coil domain of mammalian KIF1A does not form stable dimers without cargo binding (32), and we therefore stabilized human KIF1A dimers using a leucine zipper domain (9, 13). KIF1A homodimers and heterodimers were purified by the following procedure. One motor was fused with the leucine zipper and His tag, and the other was fused with the leucine zipper, a red fluorescent protein (mScarlet-I), and Strep-tag. The two constructs were coexpressed in BL21(DE3). Tandem affinity purification using TALON resin (Takara Bio Inc.) and Streptactin-XT resin (IBA Lifesciences) was performed. Eluted fractions were further separated by an NGC chromatography system (Bio-Rad) equipped with a Superdex 200 Increase 10/300 GL column (Cytiva).

### TIRF Single-Molecule Motility Assays

TIRF assays were performed as described (11, 16). Due to the highly processive nature of truncated KIF1A dimers, which often reach the end of the microtubule during experiments, we used SRP90 assay buffer [90 mM Hepes (pH 7.6), 50 mM KCH3COO, 2 mM Mg(CH3COO)2, 1 mM EGTA, 10% glycerol, biotin–bovine serum albumin (BSA) (0.1 mg/ml), K-casein (0.2 mg/ml), 0.5% Pluronic F-127] with an additional 50 mM KCH3COO in all experiments. An ECLIPSE Ti2-E microscope equipped with a CFI Apochromat TIRF 100XC oil objective lens, an Andor iXion life 897 camera, and a Ti2-LAPP illumination system (Nikon) were used to observe single-molecule motility. NIS-Elements AR software version 5.2 (Nikon) was used to control the system.

### Data Analysis

The run length and run time distributions of the homodimer and heterodimer are expected to be exponential according to our models. However, experimental limitations prevent us from measuring complete exponential distributions (33). It is impossible to include run lengths or run times shorter than the spatial or time resolution, respectively. To take account of these missing short events, we used least-squares fitting of the cumulative distribution function with a cutoff parameter (33). The experimental run length and run time data were fitted by

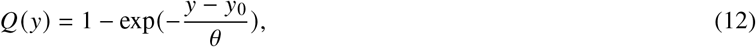

where *y* stands for *l* or *t*, and *y*_0_ and *θ* are fitting parameters.

Another problem is that KIF1A has a long run length but microtubule filaments have a limited length, resulting in a significant fraction of KIF1A motors reaching the microtubule end (22–24). Moreover, the acquisition of the image stack is also finished while motors with long run time are still moving along the filament. These limitations prevent us from observing some motors detaching normally from microtubules. To address these issues, in this study, we used microtubules longer than 50 *μ*m, which significantly exceeds the motor’s mean run length. We captured videos for 100 seconds, which is much longer than the mean run time. Additionally, the mean value *θ* obtained by least-squares fittings of Eq. 12 to the experimental data for the run length or run time was corrected by using a statistical model (25). The corrected value *θ′* is described as follows:

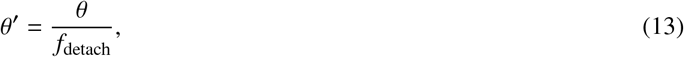

where *f*_detach_ is the fraction of motors observed detaching from microtubules. This model is based on likelihood estimation and can be applied generally for processes that generate exponential distributions (25).

The errors in the parameters obtained from these procedures can be estimated using bootstrapping (33, 34). We randomly selected measurements with replacement and divided them into two categories based on whether motor detachment was observed. We then evaluated these selected data using Eqs. 12 and 13, and obtained *θ*^′^. We repeated this procedure 10000 times to create a bootstrapping distribution. The mean of this distribution corresponds to the actual result, while the standard deviation represents the statistical error that can be expected from random sampling. This statistical error accounts for the variability in the data that would be observed if the experiment were repeated multiple times.

## RESULTS

### Analysis of KIF1A homodimers

We introduced the following KAND mutations into the *Kif1a* cDNA: KIF1A(V8M), KIF1A(R254Q), KIF1A(T258M), and KIF1A(R350G) (Fig. S4A), which have been reported as processive mutant motors (12, 14, 16). Using a single-molecule assay, we also observed robust processive movements of all these homodimeric KIF1A (Fig. S5A). The motor’s run length, run time, and velocity were measured (Fig. S5 B-D). The mean run length and run time were determined by least-squares fittings of Eq. 12 to each data (Fig. S5 B and C). Since these mean run length *λ* and run time *τ*_run_ are underestimated due to the finite microtubule lengths and observation times, as described above, each parameter was corrected using Eq. 13. To determine the detachment rate from the microtubule *k*_0d_ of each motor, we used the ADP state as a proxy for the vulnerable one-head bound state (25, 29). Single-molecule binding durations, dwell time, were measured in the saturating ADP concentrations (2 mM) (Fig. S5E). In the presence of ADP, the dwell time is theoretically expected to be distributed exponentially as *k*_0d_ exp (− *k*_0d_*t*) since the detachment in this case is a single rate-limiting process. To account for dwell times in the ADP solution which are shorter than the time resolution or interrupted by the end of the observation time, we used Eqs. 12 and 13 and obtained the corrected mean dwell time *τ*_ADP_ (= 1/ *k*_0d_) of each motor. Substituting the value of *k*_0d_ into Eqs. 3 and 5, we obtained the value of *k*_0_ and *τ*_step_. If *k*_0d_ ≪ *k*_0_, the lifetime of state 0 was approximately obtained as *τ*_0_ ≈ 1/*k*_0_. The observed and calculated parameters for each homodimer are listed in Table 1.

**Table 1:**
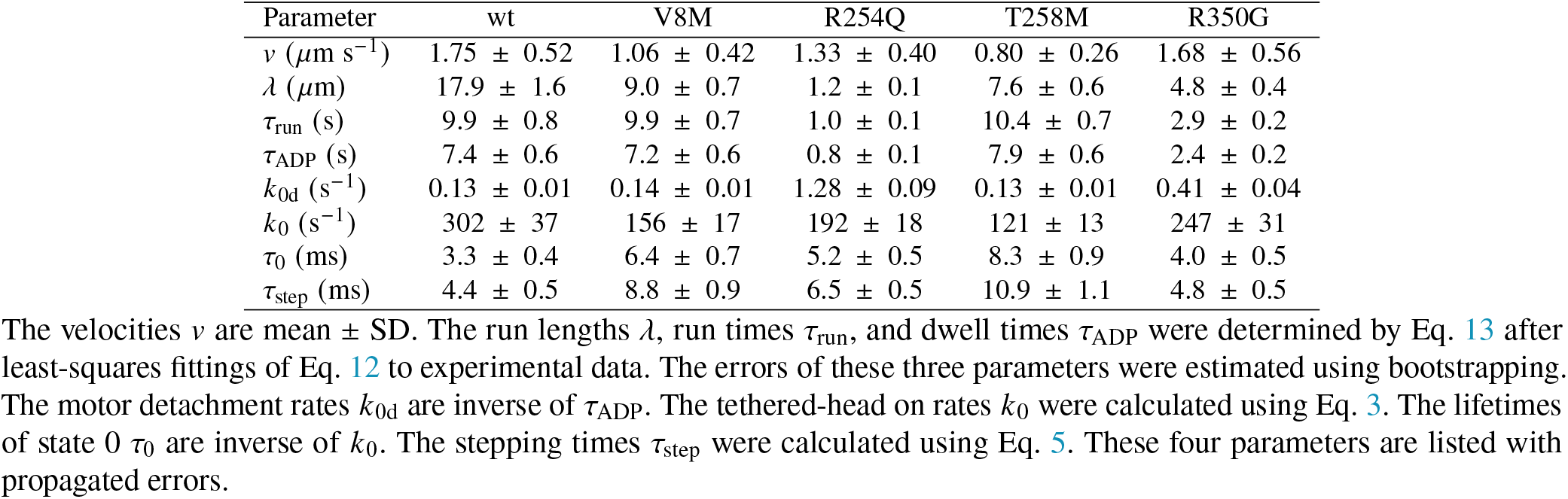
Parameters of homodimeric KIF1A obtained by single-molecule assay and calculation

It has been uncovered that the tethered-head attachment, ruled by *k*_0_, is the rate-limiting step in the KIF1A stepping cycle, indicating that KIF1A spends most of its hydrolysis cycle in the vulnerable one-head-bound state (25). Our analysis was consistent with this finding: *τ*_0_ is close to *τ*_step_ in Table 1. Therefore, the decrease in *k*_0_ significantly contributes to the reduced motor speed. Moreover, it also reduces the run length, as shown in Eq. 3, since it prolongs the duration in the weakly one-head-bound state (35). In this study, we found that all KAND mutants observed in this study shared the same rate-limiting step as wild-type KIF1A, and some of them exhibited slower *k*_0_. The V8M and T258M mutations led to a decrease in *k*_0_, resulting in slower velocity and shorter run length compared to wild-type KIF1A (Fig S5 B and D and Table 1). On the other hand, the R350G mutation reduced run length and run time due to an increase in the detachment rate from the microtubule *k*_0d_ (Fig S5 B and C and Table 1). Specifically, the R254Q mutation reduced *k*_0_ and increased *k*_0d_, leading to a decrease in all observed parameters (the velocity, run length, and run time) (Fig S5 B-D and Table 1). Based on our experiments and calculations, it appears that abnormalities in the weak one-head-bound state primarily contribute to motor performance abnormalities in certain KAND mutants.

### Analysis of KIF1A heterodimers

The single-molecule assays for the KIF1A heterodimers were conducted under the same conditions as the KIF1A homodimers. The robust processive movements were observed in all of the heterodimeric KIF1A; wt/V8M, wt/R254Q, wt/T258M, wt/R350G, V8M/R254Q, V8M/T258M, T258M/R254Q, T258M/R350G, R350G/V8M and R350G/R254Q (Fig. S6A). We observed all run length and run time distributions of the KIF1A heterodimers were adequately fit by a single exponential function (Fig. S6 B and C), which is consistent with our theory, as shown in Eqs. 7 and 9. Therefore, we determined the mean run length and run time by fitting Eq. 12 to each data using least-squares regression. To obtain the true mean run length and run time, which are not underestimated due to finite microtubule lengths and observing times, we applied the correction methods described in Eq. 13. The corrected mean run length, corrected mean run time, and mean observed velocity of each heterodimer are listed in Table 2.

**Table 2:** Motility parameters of heterodimeric KIF1A obtained by single-molecule assay

| Motor | Velocity ($\mu\text{m s}^{-1}$) | Run length ($\mu\text{m}$) | Run time (s) |
| --- | --- | --- | --- |
| wt/V8M | $1.38 \pm 0.62$ | $10.2 \pm 0.7$ | $8.5 \pm 0.6$ |
| wt/R254Q | $1.35 \pm 0.38$ | $4.3 \pm 0.4$ | $3.1 \pm 0.3$ |
| wt/T258M | $1.08 \pm 0.40$ | $10.6 \pm 0.8$ | $9.7 \pm 0.7$ |
| wt/R350G | $1.61 \pm 0.49$ | $7.8 \pm 0.6$ | $5.1 \pm 0.4$ |
| V8M/R254Q | $1.12 \pm 0.33$ | $3.5 \pm 0.2$ | $3.5 \pm 0.2$ |
| V8M/T258M | $0.90 \pm 0.27$ | $9.6 \pm 0.7$ | $11.5 \pm 0.8$ |
| T258M/R254Q | $0.98 \pm 0.25$ | $3.8 \pm 0.3$ | $4.2 \pm 0.3$ |
| T258M/R350G | $1.19 \pm 0.44$ | $7.0 \pm 0.7$ | $6.3 \pm 0.6$ |
| R350G/V8M | $1.31 \pm 0.50$ | $8.2 \pm 0.6$ | $6.9 \pm 0.4$ |
| R350G/R254Q | $1.59 \pm 0.41$ | $3.0 \pm 0.2$ | $2.0 \pm 0.1$ |

The velocities are mean ± SD. The run lengths and run times were determined by Eq. 13 after least-squares fittings of Eq. 12 to experimental data. The errors were estimated using bootstrapping.

### Determination of which head rules rate constants for heterodimer model

We initially assumed the independent head model, in which the stepping and detachment rates from the microtubule of each head in the heterodimer are identical to those in the respective homodimers. To describe the motion of the heterodimeric motors using the rate constants of the homodimers listed in Table 1, we needed to determine which head rules each reaction in the stepping cycle for the KIF1A heterodimer shown in Fig. 2. For example, should the rate *l*_0_ be identified with *k*_0_ of homodimer A or B? It is not fully understood whether this rate of tethered-head binding is ruled by the microtubule-bound head or by the tethered head itself. To investigate this uncertainty, we developed two models, model I and model II, representing the former and latter possibilities, respectively. Subsequently, we compared the predictions of these two models for the KIF1A heterodimers to determine which model better aligns with the experiment. The rate constants *l*_0_ and *l*_5_ for each model were defined as

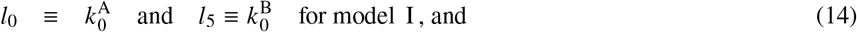

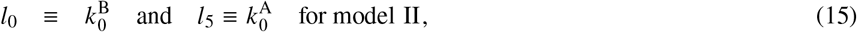

where symbols *k* ^*X*^ indicate the rate constants of head *X* obtained by analyzing homodimer *X* (*X* = A or B). The other reactions were assumed to be ruled by the head in which the corresponding reactions occur, as shown in Fig. 3A. Substituting the rate constants of the homodimers into Eqs. 8, 10, and 11, the mean run length, run time, and velocity of each model can be expressed as follows:

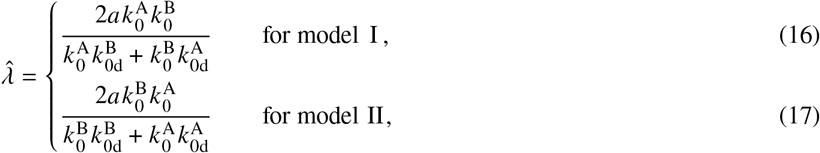

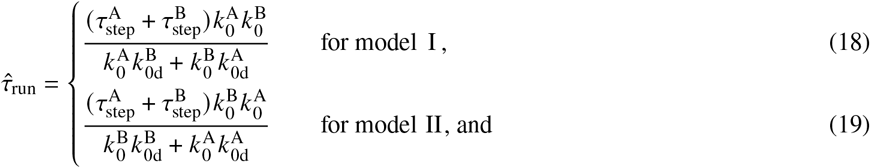

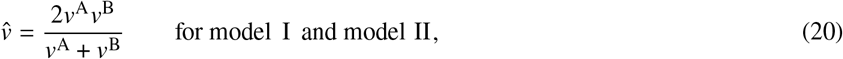

where 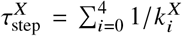 and *v*^*X*^ is the mean velocity of homodimer *X*. The run length and run time are governed by a competition between the rate of the tethered-head attachment to the next binding site, leading to a step, and the rate of premature release of the microtubule-bound head, resulting in dissociation (35). Due to differences in the degree of this competition between the two models, they predict different mean run lengths and run times. However, their predicted velocities are the same because the motor takes the same amount of time to complete two forward steps in both models.

**Figure 3:**
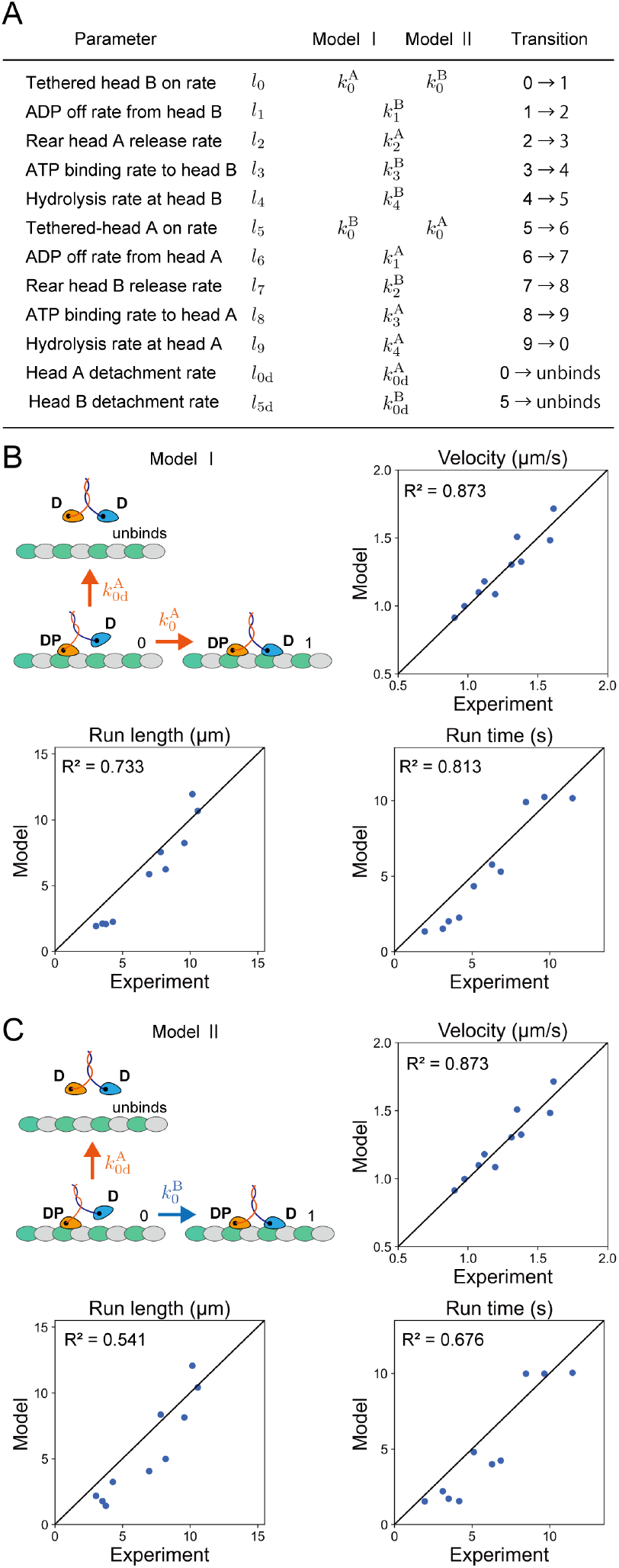
Comparison between two types of independent head models. (A) Table of rate constants for each model. (B) and (C) Correlation between experimental and predicted values. The run length, run time, and velocity of the heterodimeric KIF1A are predicted by (A) model I, in which the tethered-head attachment rate is ruled by the microtubule-bound-head and (B) model II, in which the tethered-head attachment rate is ruled by the tethered-head. The diagrams depict the transitions of each model from state 0, as an example, which is involved in the kinetic race between the tethered head attachment rate and microtubule detachment rate. The experimental and predicted values are plotted with the coefficient of determination *R*^2^ score values.

The two models were compared using the coefficient of determination (*R*^2^ score). The model I achieved a higher *R*^2^ score than the model II when comparing the experimental and predicted values for the run lengths and run times (Fig. 3B and C).

The predicted velocities of the model I, which are the same as those of the model II, also showed good agreement with the experimental data (Fig. 3B and C). Based on these results, we concluded that the rate of tethered-head attachment is determined by the properties of the microtubule-bound head. For the remainder of this article, we primarily use the model I.

### Detailed comparison between experimental and model values

Experimental data demonstrated that when there was a disparity in the performance of two motors, the motility of the heterodimer composed of them tended to be approximately intermediate or closer to the lower value (Fig. 4A-C). The independent head model I was able to roughly capture this trend using the parameters of the homodimers (Fig. 4A-C). However, we consistently observed discrepancies in the run length between the experimental data and the model for every R254Q-related heterodimer. While our model calculated that the run lengths of R254Q-related heterodimers (2.2 ± 0.1 *μ*m, 2.1 ± 0.1 *μ*m, 2.1 ± 0.1 *μ*m and 1.9 ± 0.1 *μ*m, for wt/R254Q, V8M/R254Q, T258M/R254Q, and R350G/R254Q respectively) were much closer to the value of the homodimer composed of R254Q (1.2 ± 0.1 *μ*m), the experimental values for these heterodimers (4.3 ± 0.4 *μ*m, 3.5 ± 0.2 *μ*m, 3.8 ± 0.3 *μ*m and 3.0 ± 0.2 *μ*m, for wt/R254Q, V8M/R254Q, T258M/R254Q, and R350G/R254Q respectively) were 1.5–2.0 times larger than predicted (Fig. 4A). Similarly, their mean run times were underestimated by the independent head model (Fig. 4B). In this model, we assumed that all reactions were ruled by either of the heads, but our results indicated that some reactions are influenced by both heads rather than just one. Since the independent head model successfully explained the velocity of heterodimers (Fig. 4C), it was reasonable to consider that the rate constants in the step direction were accurate but the detachment rate was not adequately captured in this model.

**Figure 4:**
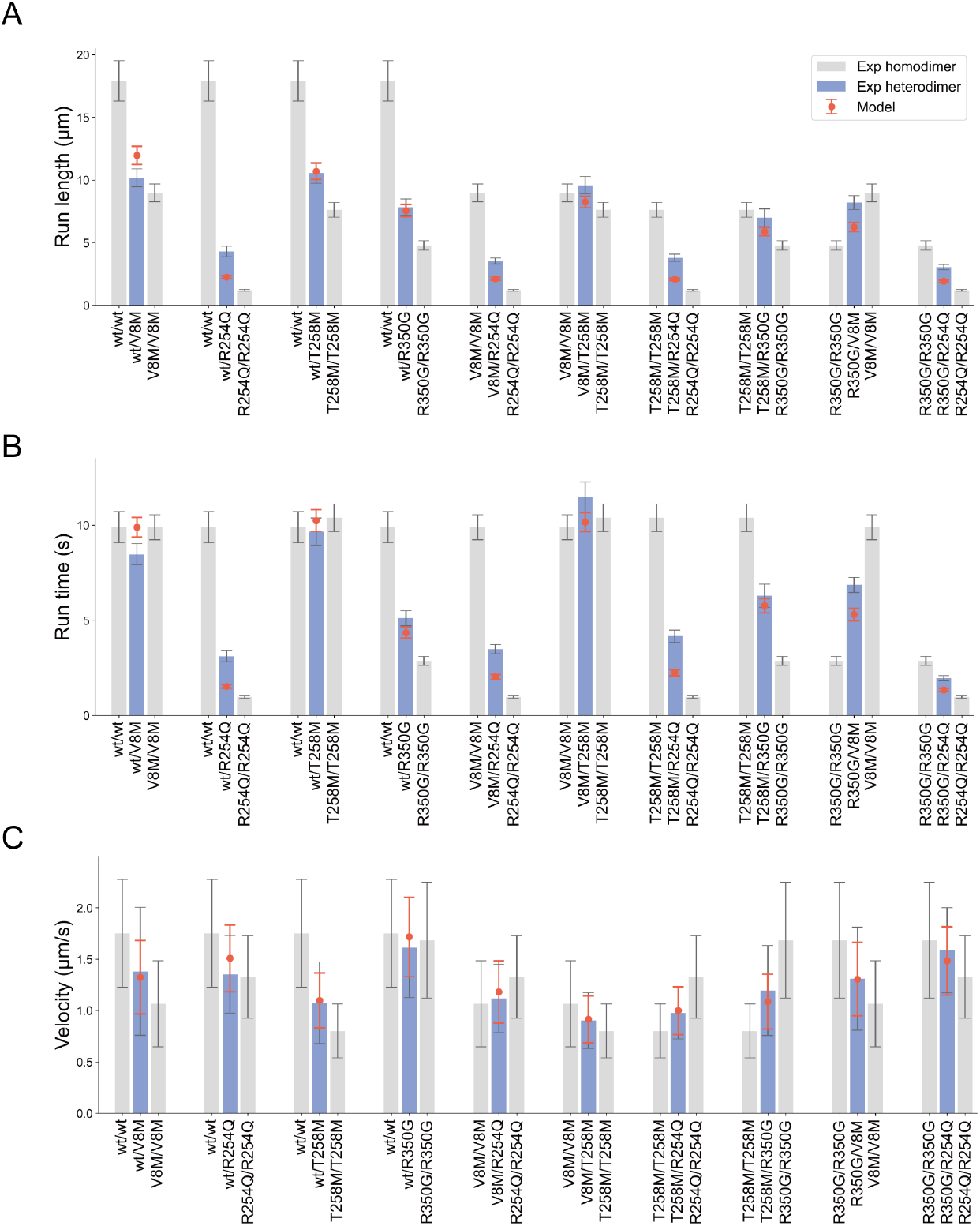
Detailed comparison of experimental and predicted values in (A) run length, (B) run time, and (C) velocity. Gray bars and blue bars show the experimental values of homodimers and heterodimers, respectively. Orange plots show the predicted values of heterodimers by the independent head model I. The experimental mean run length and run time were determined by Eq. 13 after least-squares fittings of Eq. 12 to data. The error bars represent the mean ± bootstrapping error. The predicted mean run length and run time of the heterodimer were calculated using Eqs. 16 and 18, respectively. The error bars represent the mean ± propagated error. The experimental velocity is mean ± SD. The predicted mean velocity was calculated using Eq. 20 and the error bars represent the mean ± propagated error.

### Introduction of tethered head and microtubule interactions

To improve our model, we incorporated the interactions between the tethered head and the microtubule by modifying the detachment rates, *l*_0d_ and *l*_5d_. Previous studies have suggested that the tethered head of kinesin-1 exhibits highly diffusive movement on the microtubule surface before binding to the forward binding site (36, 37). Now, we constructed a model in which the tethered head interacts with the microtubule while undergoing diffusive movement on the microtubule surface, as shown in Fig. 5A. We decomposed state 0 into two sub-states: 0′ and 0′′. In state 0′, there are no interactions between the tethered head and the microtubule, while in state 0′′, there are interactions between them. After ATP hydrolysis in the microtubule-bound head in state 4, the motor transitions to state 0′ (transition 4 → 0′). The transitions between states 0′ and 0′′ are reversible and governed by rate constants *k*_0_′ and *k*_0_′′. We assumed that detachment from the microtubule occurs only in state 0′ with a rate of *k*_0_′_d_ (transition 0′→ unbinds). In our model, the attachment of the tethered head to the microtubule is ruled by the microtubule-bound head; therefore, both transitions 0′→ 1 and 0′′→ 1 occur at the same rate as the conventional *k*_0_. It is possible that *k*_0_′ and *k*_0_′′ are orders of magnitude faster than the other rate constants in the stepping cycle shown in Fig. 1 because the former transitions are primarily driven by thermal fluctuations. By considering these conditions, we were able to introduce the interactions between the tethered head and the microtubule without adding new reaction pathways to the stepping cycle shown in Fig. 1 (see Section S3 of the Supporting Materials and Methods and Fig. S1C). The conventional detachment rate *k*_0d_ can be decomposed as

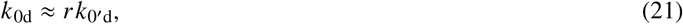

where *r* defined by

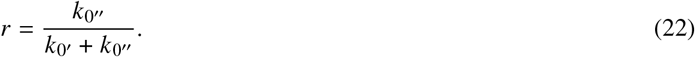

represents the proportion of time the motor spends in the state 0′ after the transition between the states 0′ and 0′′ has immediately reached equilibrium. Note that according to Eqs. 21 and 22, *r* and *k*_0_′_d_ are associated with the tethered head and the microtubule-bound head, respectively.

**Figure 5:**
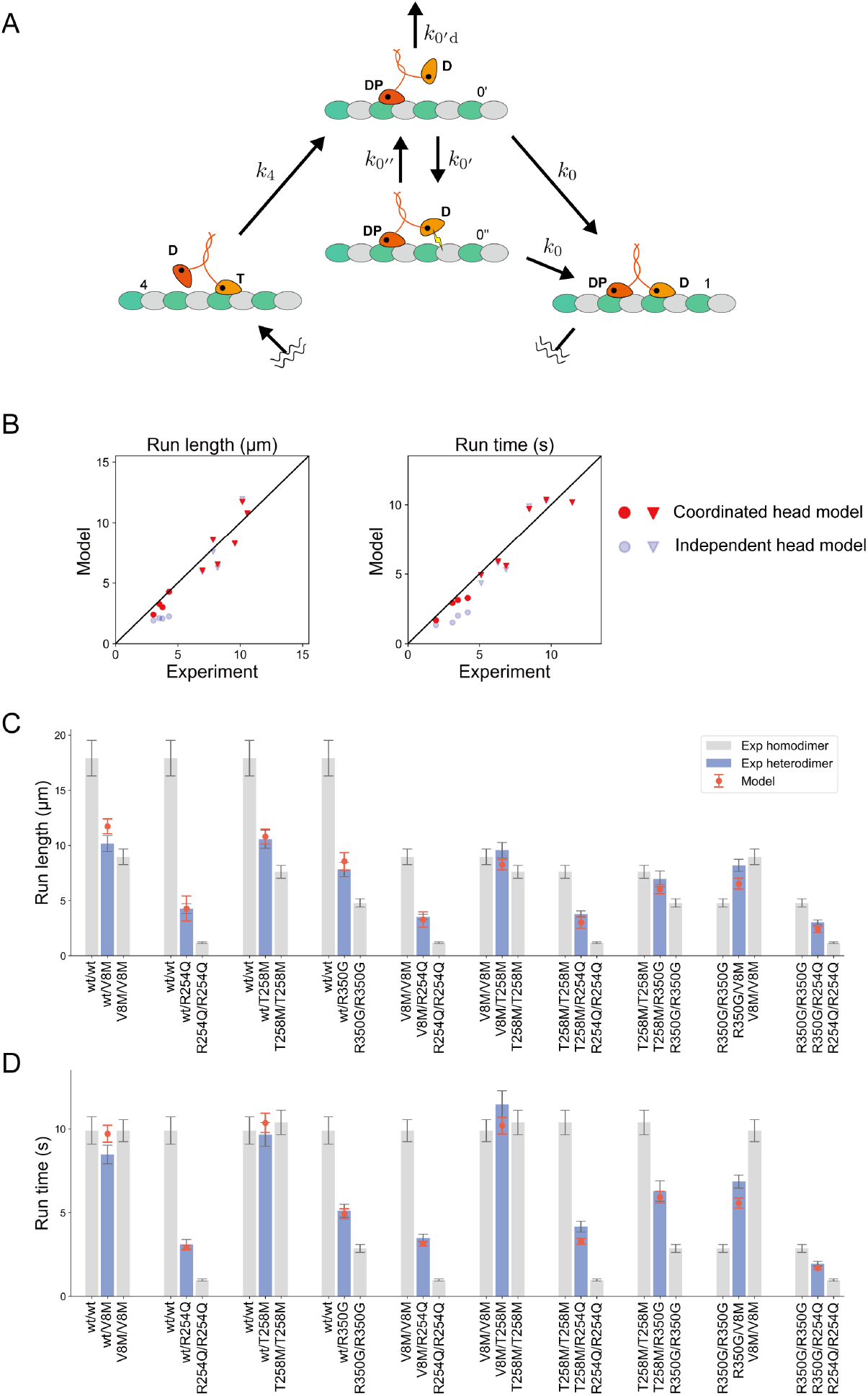
The coordinated head model in which tethered-head interacts with the microtubule. (A) Modified stepping cycle for homodimeric KIF1A. A model consists of six states and states 1∼4 are the same states as in the model shown in Fig. 1. States 2 and 3 are not indicated in a diagram. Conventional state 0 is decomposed into two sub-states: 0′ and 0′′. State 0′′ has interactions between the tethered head and the microtubule, indicated by a yellow lightning symbol, while state 0′ does not involve such interactions. After ATP is hydrolyzed in state 4, the motor transitions to state 0′. Transitions between 0′ and 0′′, governed by *k*0′ and *k*0′′, are reversible. From state 0′ and 0′′, the motor transitions to state 1 at *k*_0_. Otherwise, the motor detaches from the microtubule at *k*_0_′_d_ in state 0′ and terminates the processive run. (B) Correlation between experimental and predicted values. The experimental values are compared with the values of the coordinated head model, plotted in red, and the independent head model, plotted in blue. Circles represent the results for R254Q-related heterodimers, while triangles represent the results for other heterodimers. (C and D) Detailed comparison between experimental values and predicted values by the coordinated head model for (C) run length and (D) run time. Gray bars and blue bars represent the same experimental values of homodimers and heterodimers, respectively, as shown in Fig. 4. Orange plots represent the predicted values of heterodimers by the coordinated head model. The predicted mean run length and run time of the heterodimers were calculated using Eqs. 24 and 25, respectively. The error bars represent the mean ± propagated error.

We made this modification on the model for the KIF1A heterodimers, which we refer to as the coordinated head model in this article. The rates of dissociation from the microtubule in states 0 and 5 were defined as follows:

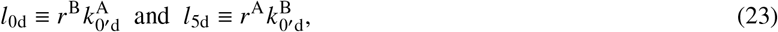

where 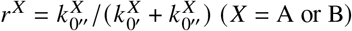. The detachment of the motor in each vulnerable one-head bound state depends on the properties of the two different heads. The rate constants other than *l*_0d_ and *l*_5d_ are the same as those in the independent head model I shown in Fig. 3A. The mean run length and run time in the coordinated head model are obtained by substituting Eq. 23 into Eqs. 16 and 18, respectively:

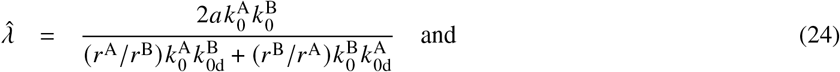

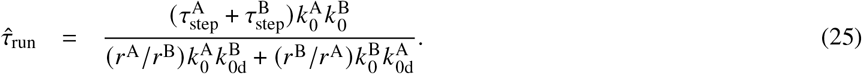

As the rate constants in the step direction are the same as those of the independent head model, the modification does not affect the predicted motor velocity. Although it seemed difficult to experimentally determine the value of *r* for each motor, the ratio *r*^A^ / *r*^B^ associated with motors A and B could be estimated from the experimental value of *r* for the heterodimer composed of motors A and B and Eq. 24 (Fig. S7). The R254Q motor was chosen as a reference motor, and the ratios *r*^A^ / *r*^R254Q^ with various A were estimated to obtain *r*^wt^ / *r*^R254Q^ = 0.39, *r*^V8M^ / *r*^R254Q^ = 0.37, *r*^T258M^ / *r*^R254Q^ = 0.40, and *r*^R350G^ / *r*^R254Q^ = 0.50. Note that these values are not significantly different from each other.

We used the parameters of homodimeric motors listed in Table 1 and the values of *r*^A^ / *r*^R254Q^ to predict the mean run lengths and run times of the KIF1A heterodimers. The predicted and experimental values of these quantities were compared in Fig. 5B–D. Comparing Figs. 5C and 5D with Figs. 4A and 4B, respectively, we see that the agreement between the theory and experiment was improved by the modification of the model for the R254Q-relate heterodimers. For the other heterodimers, the theory remained consistent with experimental data. This is because *r*^A^ / *r*^R254Q^ had similar values, as noted above, and the predicted run lengths and run times of these heterodimers were not significantly altered by our modification. The incorporation of interactions between the tethered head and the microtubule allowed us to better characterize the motility of the KIF1A heterodimers.

## DISCUSSION

### Practicality of our models

According to our theory, the run length and run time distributions of the highly processive heterodimeric motor can be approximately described by a single exponential function. However, each distribution near the starting point (the zero of run length or run time) is influenced by which head binds to the microtubule first and does not follow a single exponential distribution. Since this behavior only appears in a narrow region of their distributions, this effect cannot be detected with the time and spatial resolutions of TIRF microscopy (in our case, 0.10 s and 0.16 *μ*m, respectively) (Fig. S3). Therefore, their experimental distributions obtained through TIRF assays can be adequately fitted by a single exponential distribution.

We developed two models, the independent head model and the coordinated head model, to describe the motion of heterodimeric KIF1A by using parameters obtained from homodimers (Fig. S8). The independent head model, which considers the attachment of the tethered head being ruled by the microtubule-bound head, provided approximate predictions of the motility of heterodimeric KIF1A (Fig. 4). This model allows us to gain insight into the motility parameters of disease-associated KIF1A heterodimers by separately purifying and analyzing each homodimer. The model can be simplified by using only the run lengths and run times of the homodimeric motors instead of their detailed rate constants (see Section S4 of the Supporting Materials and Method). This simplification allows experimenters to readily utilize the model without conducting experiments in ADP solutions. On the other hand, the coordinated head model is not suitable for predicting the motility of heterodimers using homodimer parameters since it relies on parameters obtained from analyzing a different heterodimer to describe the motility of the specific heterodimer of interest. However, such model refinement can better characterize the motor’s movement and provide a more plausible walking mechanism (Fig. 5A). Researchers can choose the appropriate model depending on their specific purpose.

### Limitation of our models

The models developed for homodimers and heterodimers assume that the motor moves forward processively along filaments in a hand-over-hand manner. Therefore, they cannot be applied to heterodimeric motors composed of processive and non-processive motors. It has been demonstrated that heterodimerization with a processive partner can generate processivity when one motor is not processive as a homodimer, as observed in the study of KIF1A (16) and other motors (38, 39). Most non-processive homodimeric KIF1A exhibits one-dimensional Brownian motion on the microtubule (9, 16, 29). Therefore, it is necessary to consider diffuse motion and additional reaction pathways in the motor stepping cycle to describe the processive behavior of heterodimers composed of processive and non-processive motors.

To assess the applicability of our models to other kinesin motors, it is necessary to validate their suitability. Kinesin-1, which has the same stepping cycle as KIF1A (36, 40), is a potential candidate for applying our independent head model because it is likely that each wild-type and mutant head of heterodimeric kinesin-1 independently and sequentially steps along the microtubule (19). However, applying our models to heterodimeric kinesin-2, particularly KIF3AC, presents a challenge despite its similar stepping cycle to KIF1A (41). KIF3AC exhibits a longer run length compared to both KIF3AA and KIF3CC, and its speed is much faster than expected based on the very slow rate of the KIF3CC (31, 42). Additionally, KIF3AC does not demonstrate alternating dwell times corresponding to the KIF3A head and KIF3C head (31). It has been discovered that KIF3A significantly alters the properties of KIF3C through heterodimerization (21, 43); therefore, modeling the behavior of KIF3AC using parameters from KIF3AA and KIF3CC would require making certain assumptions.

Our models are intentionally designed to be simple, and we believe it would be efficient to build customized models based on them to accurately capture the unique behavior exhibited by these specific motors.

### Cooperation between KIF1A heads

The tethered head attachment is identified as a rate-limiting step in the KIF1A hydrolysis cycle (Table 1) (25). We observed that all of the homodimer mutants examined in this study exhibit the same rate-limiting step as wild-type KIF1A (Table 1). Therefore, if the tethered head attachment is ruled by either head, the velocity of the KIF1A heterodimer is primarily governed by two types of tethered head attachment rates or the significantly slower one. This hypothesis was confirmed by the good agreement between the independent head model and the velocity of KIF1A heterodimers (Fig. 4C).

Our models further determined that the attachment of the tethered head to the microtubule is likely governed by the microtubule-bound head rather than the tethered head, which is consistent with the previous experiments (25, 29). The full docking of the neck linker in the microtubule-bound head is possibly essential for the attachment of the tethered head (25), and our analysis of mutant homodimers also supports this conclusion. We revealed that the KIF1A(V8M), KIF1A(R254Q), and KIF1A(T258M) have slower tethered head attachment rates compared to wild-type KIF1A according to our calculations. V8 is located at the *β*1 sheet, while R254 and T258 are within the L11 loop. Since both of these regions play a crucial role in nucleotide-dependent neck-linker docking, it is likely that mutations in these regions would affect the speed of the neck linker docking (14, 44, 45).

On the other hand, with regard to the association between the KIF1A dimer and the microtubule, both the microtubule-bound head and the tethered head may play critical roles. Our coordinated head model suggests that the tethered head interacts with the microtubule, thereby suppressing the dissociation of the microtubule-bound head. Notably, KIF1A possesses a lysine-rich insertion in the L12 loop known as “K-loop”, which allows the monomeric KIF1A head to undergo biased Brownian motion on the microtubule without fully detaching from it. The K-loop is also essential for the superprocessive movement of the KIF1A dimer under near-physiological ionic strength conditions (29, 46, 47). One potential factor contributing to the interaction between the tethered head and the microtubule is the presence of the K-loop.

Our models provide insights into the movement of heterodimeric KIF1A and highlight the potential roles of both the microtubule-bound head and the tethered head in governing the fast velocity and superprocessivity of the KIF1A dimer. These findings contribute to our understanding of the intricate interplay between KIF1A and the microtubule. It is anticipated that applying our models to other molecular motors, either as they are or with some modifications, will help uncover the molecular mechanisms underlying motor function.

## AUTHOR CONTRIBUTIONS

T.K., K.S. and S.N. designed research; T.K and S.N. performed experiments; T.K. and K.S. performed calculations; T.K analyzed data; T.K., K.S. and S.N. wrote the paper.

## ACKNOWLEDGMENTS

Generous support from the FRIS CoRE, which is a shared research environment. T.K. was supported by a Grant-in-Aid of Tohoku University, Division for Interdisciplinary Advanced Research and Education. SN was supported by JSPS KAKENHI (grants nos. 23H02472 and 22H05523).

