## Supplemental Information for "Modeling the motion of disease-associated KIF1A heterodimers"

### SUPPORTING METHODS

### S1 Model for homodimeric motor

We consider the stepping cycle for KIF1A homodimers as shown in Fig. 1 of the main text, which consists of five states. A pathway represented by a solid line with symbol $k_i$ indicates the reaction in a forward step. Irreversible detachment of the motor in state 0 from the track is represented by the vertical arrow with the symbol $k_{0\text{d}}$. We are interested in studying run length $\lambda$ and run time $\tau_{\text{run}}$ of the homodimeric motor.

#### S1-1 Run length of homodimeric motor

The probability density function for the run length can be calculated as follows. The probability of the motor detaching per step $p_{0\text{d}}$ is given as,

$$p_{0\text{d}} = \frac{k_{0\text{d}}}{k_0 + k_{0\text{d}}}. \tag{S1}$$

Note that the motor is highly processive if $p_{0\text{d}} \ll 1$ (i.e., $k_{0\text{d}} \ll k_0$), because the detachment hardly occurs before taking the next step under this condition. The probability that the motor repeats steps $n$ times and detaches without taking the next step is as follows:

$$\pi_n = p_{0\text{d}}(1 - p_{0\text{d}})^n = p_{0\text{d}} \exp(-bn), \tag{S2}$$

where a positive constant $b$ is defined by

$$b \equiv -\ln(1 - p_{0\text{d}}). \tag{S3}$$

Suppose that we measure the run length by a device with spatial resolution of $\delta$ when the motor detaches after taking $n$ steps with a step size of $a$. Then, the probability density distribution for the measured run length $l$ will be given by the normal distribution:

$$R_n(l) = \frac{1}{\sqrt{2\pi}\delta} \exp\left[-\frac{(l - na)^2}{2\delta^2}\right]. \tag{S4}$$

Now, the run length distribution obtained by measuring a large number of run lengths can be expressed as follows:

$$P_{\text{RL}}(l) = \sum_{n=0}^{\infty} \pi_n R_n(l). \tag{S5}$$

To derive approximate expressions for the infinite sum in Eq. S5 under certain conditions, we introduce symbols $\lambda$, $\alpha$, $\beta$, and $c$ defined by

$$\lambda \equiv \frac{a}{b}, \quad \alpha \equiv \frac{a^2}{2\delta^2}, \quad \beta \equiv \frac{l}{a} - \frac{\delta^2}{a\lambda}, \quad c \equiv \frac{1}{a} \exp\left[\frac{1}{2}\left(\frac{\delta}{\lambda}\right)^2\right] \tag{S6}$$

and function $f(x)$ defined by

$$f(x) \equiv \sqrt{\frac{\alpha}{\pi}} \exp[-\alpha(x - \beta)^2]. \tag{S7}$$

Now, substituting these expressions into Eq. S5, we obtain

$$P_{\text{RL}}(l) = c p_{0\text{d}} \exp\left(-\frac{l}{\lambda}\right) \sum_{n=0}^{\infty} f(n). \tag{S8}$$

According to the Poisson summation formula, the sum in Eq. S8 can be rewritten as

$$\sum_{n=0}^{\infty} f(n) = \int_0^{\infty} f(x) \left[1 + 2\sum_{h=1}^{\infty} \cos(2\pi hx)\right] dx - \frac{1}{2} f(0). \tag{S9}$$

In what follows we assume that $p_{0d} \ll 1$, i.e. the motor is highly processive. In this case, $b$ defined in Eq. S3 can be approximated as $b \approx p_{0d}$, and $\lambda$ defined in Eq. S6 as

$$\lambda \approx \frac{a}{p_{0d}} \approx \frac{a k_0}{k_{0d}}. \tag{S10}$$

Note that $1/p_{0d}$ is approximately the average number of steps before detachment, i.e. $\sum_{n=0}^{\infty} n\pi_n$, in this case. Therefore, $\lambda$ represents the approximate average run length. We also assume that the average run length is much larger than the spatial resolution of the device, i.e. $\lambda \gg \delta$; otherwise, we cannot obtain reliable data experimentally. Under these conditions $c$ defined in Eq. S6 can be approximated as $c = 1/p_{0d}\lambda$, and Eq. S8 as

$$P_{\mathrm{RL}}(l) \approx \frac{1}{\lambda} \exp\left(-\frac{l}{\lambda}\right) \sum_{n=0}^{\infty} f(n). \tag{S11}$$

The probability density $P_{\mathrm{RL}}(l)$ can be further simplified, if the spacial resolution $\delta$ is larger than the step size $a$. Under this condition, we have $\alpha \ll 1$ according to the second equation in Eq. S6. This implies that function $f(x)$ defined in Eq. S7 hardly changes its value when $x$ varies over an interval of range $1/h$ ($h = 1, 2, \dots$), the period of $\cos(2\pi h x)$ in Eq. S9. Therefore, the contribution from the second term in the square brackets in this equation to the integral should be negligibly small, which will be ignored in the following calculations. The remaining integral in Eq. S9 can be expressed in terms of the complementary error function defined by

$$\mathrm{erfc}(x) = \frac{2}{\sqrt{\pi}} \int_x^{\infty} \exp(-z^2) dz,$$

and Eq. S11 is now approximated as

$$P_{\mathrm{RL}}(l) \approx \frac{1}{\lambda} \exp\left(-\frac{l}{\lambda}\right) \left[1 - \frac{1}{2}\mathrm{erfc}(\sqrt{\alpha}\beta) - \frac{1}{2}\sqrt{\frac{\alpha}{\pi}} \exp(-\alpha\beta^2)\right]. \tag{S12}$$

Remember that this approximation is valid under the conditions $\lambda \gg a$ (i.e. $p_{0d} \ll 1$), $\lambda \gg \delta$, and $\delta \gg a$.

Finally, if $l^2 \gg \delta^2$ is satisfied, then we have $\alpha\beta^2 \gg 1$ provided that $\lambda \gg \delta$, which is one of the conditions mentioned above. Under these conditions, the error function can be approximated as $\mathrm{erfc}(\sqrt{\alpha}\beta) \approx \exp(-\alpha\beta^2)/\sqrt{\pi\alpha\beta^2}$. Then, the last two terms in the square brackets in Eq. S12 are negligible compared with the first, and the following approximation is obtained:

$$P_{\mathrm{RL}}(l) \approx \frac{1}{\lambda} \exp\left(-\frac{l}{\lambda}\right). \tag{S13}$$

Eqs. S10 and S13 correspond to Eqs. 3 and 2 in the main text, respectively.

#### S1-2 Run time of homodimeric motor

To obtain the run time distribution of the homodimeric motor, we introduce the following rate constant, which approximates the transitions $1 \to 2 \to 3 \to 4 \to 0$:

$$K_1 = \left(\frac{1}{k_1} + \frac{1}{k_2} + \frac{1}{k_3} + \frac{1}{k_4}\right)^{-1}. \tag{S14}$$

We assume a two-state model governed by $k_0$, $k_{0d}$, and $K_1$ shown in Fig. S1A, in which the motor starts running in state 0 at time $t = 0$. Let $Q_i$ be the probability that the motor is in state $i$ ($i = 0, 1$) on the microtubule and we have the following master equations:

$$\begin{aligned} \frac{dQ_0(t)}{dt} &= K_1 Q_1(t) - (k_0 + k_{0d}) Q_0(t), \\ \frac{dQ_1(t)}{dt} &= k_0 Q_0(t) - K_1 Q_1(t). \end{aligned} \tag{S15}$$

The solutions to these coupled equations with the initial conditions $Q_0(0) = 1$ and $Q_1(0) = 0$ are as follows:

$$\begin{aligned} Q_0(t) &= \frac{\alpha_+ - K_1}{\alpha_+ - \alpha_-} \exp(-\alpha_+ t) - \frac{\alpha_- - K_1}{\alpha_+ - \alpha_-} \exp(-\alpha_- t) \\ Q_1(t) &= -\frac{k_0}{\alpha_+ - \alpha_-} \left[\exp(-\alpha_+ t) - \exp(-\alpha_- t)\right], \end{aligned} \tag{S16}$$

where

$$\alpha_\pm = \frac{1}{2}\left[k_0 + k_{0d} + K_1 \pm \sqrt{(k_0 + k_{0d} + K_1)^2 - 4k_{0d}K_1}\right]. \tag{S17}$$

The probability $Q(t)$ of the motor being on the microtubule at time $t$ can be expressed as $Q(t) = Q_0(t) + Q_1(t)$, and the run time distribution $P_{\mathrm{RT}}(t)$ is obtained as $P_{\mathrm{RT}}(t) = -dQ(t)/dt$, which results in

$$P_{\mathrm{RT}}(t) = \frac{k_{0d} - \alpha_-}{\alpha_+ - \alpha_-}\alpha_+ \exp(-\alpha_+ t) + \frac{\alpha_+ - k_{0d}}{\alpha_+ - \alpha_-}\alpha_- \exp(-\alpha_- t). \tag{S18}$$

If the motor is highly processive ($k_0, K_1 \gg k_{0d}$), $\alpha_+$ and $\alpha_-$ can be approximated as follows:

$$\alpha_+ \approx k_0 + K_1, \tag{S19}$$

$$\alpha_- \approx \frac{k_{0d}K_1}{k_0 + K_1} \tag{S20}$$

For $t \gg \alpha_+^{-1}$ ($\alpha_+^{-1} \sim 1$ ms for the homodimers used in this study; Table 1 in the main text), the run time distribution of the highly processive homodimeric motor is given as

$$P_{\mathrm{RT}}(t) \approx \alpha_- \exp(-\alpha_- t), \tag{S21}$$

and the mean run time is

$$\tau_{\mathrm{run}} = \frac{1}{\alpha_-} = \frac{\tau_{\mathrm{step}} k_0}{k_{0d}}, \tag{S22}$$

where

$$\tau_{\mathrm{step}} = \frac{1}{k_0} + \frac{1}{K_1} = \sum_{i=0}^{4} \frac{1}{k_i}. \tag{S23}$$

Eqs. S21 and S22 correspond to Eqs. 4 and 5 in the main text, respectively.

#### S1-3 Inspection of the model for homodimeric motor

For motors with a fixed step size, such as kinesin, the actual run length distribution consists of almost discrete values. However, it is impossible to observe the discrete run length distribution at our spatial resolution of 160 nm, which is greater than the step size (8 nm for KIF1A) (Figs. S2A–C). If the motor exhibits high processivity, the run length distribution can be approximately described by a single exponential function given in Eq. S13, which is commonly used (1–3), except in a narrow interval near the zero of run length (Fig. S2C).

To obtain the run time distribution, we introduced the rate constant $K_1$, which approximates the transitions $1 \longrightarrow 2 \longrightarrow 3 \longrightarrow 4 \longrightarrow 0$, as described in Eq. S14. This approximation is reasonable when one rate constant is significantly smaller than the others included in $K_1$. In the case of KIF1A, it satisfies this condition as the ADP off rate from the front head $k_1$ is at least four times smaller than $k_2$, $k_3$, and $k_4$ (2).

In contrast to the run length, the run time $t$ is continuous valued. However, the run time distribution for a highly processive motor can be described by a single-exponential function in Eq. S21, similar to the run length distribution, except in a narrow interval near $t = 0$. We have $P_{\mathrm{RT}}(0) = k_{0d}$ from Eq. S18, which is somewhat larger than $P_{\mathrm{RT}}(0) = \alpha_-$ calculated from Eq. S21; this is because we chose the weakly one-head-bound state (state 0) after ATP hydrolysis as the initial state. The deviation from the single-exponential behavior at the initial stage disappears after the relaxation time of order $\alpha_+^{-1}$. A similar transient behavior can, in principle, be observed in an actual motor, because it initiates its run from the vulnerable one-head-bound state upon binding to the microtubule from the solution (2). However, to observe this behavior in the distribution, a time resolution on the order of microseconds is required (Fig. S2D). Therefore, in TIRF assays with our time resolution of 100 ms, the run time distribution appears to follow a single exponential distribution.

### S2 Model for heterodimeric motor

We consider the heterodimer model in Fig. 2 consisting of ten states. A pathway represented by a solid line with symbol $l_i$ indicates the reaction in a forward step. Irreversible detachment of the motor in states 0 and 5 from the track is represented by the vertical arrow with symbol $l_{0d}$ and $l_{5d}$, respectively. We are interested in the run length and run time of the heterodimeric motor.

### S2-1 Run length of heterodimeric motor

The probability density function for the run length of the heterodimeric motor can be calculated as follows. The probabilities of the motor detaching in state 0 and in state 5 are given as

$$p_{0\mathrm{d}} = \frac{l_{0\mathrm{d}}}{l_0 + l_{0\mathrm{d}}}, \qquad p_{5\mathrm{d}} = \frac{l_{5\mathrm{d}}}{l_5 + l_{5\mathrm{d}}}. \tag{S24}$$

The motor is highly processive if both $p_{0\mathrm{d}} \ll 1$ and $p_{5\mathrm{d}} \ll 1$ (i.e. $l_{0\mathrm{d}} \ll l_0$ and $l_{5\mathrm{d}} \ll l_5$) are satisfied. We assume that the motor starts running in state 0. The probability $\pi_{2m}$ that the motor repeats steps $2m$ times and detaches without taking the next step and the probability $\pi_{2m+1}$ that the motor repeats steps $2m+1$ times and detaches without taking the next step can be expressed as follows:

$$\begin{aligned} \pi_{2m} &= p_{0\mathrm{d}}(1-p_{0\mathrm{d}})^m(1-p_{5\mathrm{d}})^m = p_{0\mathrm{d}}\exp(-bm), \\ \pi_{2m+1} &= p_{5\mathrm{d}}(1-p_{0\mathrm{d}})^{m+1}(1-p_{5\mathrm{d}})^m = \sqrt{\frac{1-p_{0\mathrm{d}}}{1-p_{5\mathrm{d}}}}\,p_{5\mathrm{d}}\exp\left[-b\left(m+\frac{1}{2}\right)\right], \end{aligned} \tag{S25}$$

where a positive constant $b$ is defined by

$$b \equiv -\ln(1-p_{0\mathrm{d}})(1-p_{5\mathrm{d}}). \tag{S26}$$

Using the normal distribution $R_n(l)$, Eq. S4, representing the spacial resolution of the device, the measured run length distribution of the heterodimeric motor is given as follows:

$$\hat{P}_{\mathrm{RL}}(l) = \sum_{m=0}^{\infty}\left[\pi_{2m}R_{2m}(l) + \pi_{2m+1}R_{2m+1}(l)\right]. \tag{S27}$$

To rewrite this equation, we introduce symbols $\hat{\lambda}$, $\alpha$, $\beta_0$, $\beta_5$, and $c$ defined by

$$\hat{\lambda} \equiv \frac{2a}{b}, \quad \alpha \equiv \frac{2a^2}{\delta^2}, \quad \beta_0 \equiv \frac{l}{2a} - \frac{\delta^2}{2a\hat{\lambda}}, \quad \beta_5 \equiv \beta_0 - \frac{1}{2}, \quad \text{and} \quad c \equiv \frac{1}{2a}\exp\left[\frac{1}{2}\left(\frac{\delta}{\hat{\lambda}}\right)^2\right] \tag{S28}$$

and functions $f_j(x)$ defined by

$$f_j(x) = \sqrt{\frac{\alpha}{\pi}}\exp[-\alpha(x-\beta_j)^2] \quad (j = 0, 5). \tag{S29}$$

Now, substituting these expressions into Eq. S27, we have the following:

$$\hat{P}_{\mathrm{RL}}(l) = c\,p_{0\mathrm{d}}\exp\left(-\frac{l}{\hat{\lambda}}\right)\sum_{m=0}^{\infty} f_0(m) + c\sqrt{\frac{1-p_{0\mathrm{d}}}{1-p_{5\mathrm{d}}}}\,p_{5\mathrm{d}}\exp\left(-\frac{l}{\hat{\lambda}}\right)\sum_{m=0}^{\infty} f_5(m). \tag{S30}$$

Using the Poisson summation formula, the sums in Eq. S30 can be rewritten as Eq. S9 with $f(x)$ replaced by $f_j(x)$.

In what follows we assume that the motor is highly processive, i.e. $p_{0\mathrm{d}} \ll 1$ ($l_{0\mathrm{d}} \ll l_0$) and $p_{5\mathrm{d}} \ll 1$ ($l_{5\mathrm{d}} \ll l_5$) are satisfied. In this case, $b$ defined in Eq. S26 can be approximated as $b \approx p_{0\mathrm{d}} + p_{5\mathrm{d}}$, and $\hat{\lambda}$ defined in Eq. S28 is approximated as

$$\hat{\lambda} \approx \frac{2a}{p_{0\mathrm{d}} + p_{5\mathrm{d}}} \approx \frac{2al_0l_5}{l_0l_{5\mathrm{d}} + l_5l_{0\mathrm{d}}}, \tag{S31}$$

which means the mean run length of the highly processive motor. We also assume that $\hat{\lambda}^2 \gg \delta^2$ is satisfied for the same reason as the case of homodimeric motor. Under these conditions $c$ defined in Eq. S28 is approximated as $c \approx 1/2a$ and Eq. S30 as

$$\hat{P}_{\mathrm{RL}}(l) \approx \frac{1}{2a}\exp\left(-\frac{l}{\hat{\lambda}}\right)\sum_{m=0}^{\infty}\left[p_{0\mathrm{d}}f_0(m) + p_{5\mathrm{d}}f_5(m)\right]. \tag{S32}$$

As in the case of the model for homodimeric motor, if $a^2 \ll \delta^2$ is satisfied, Eq. S32 can be simplified as

$$\hat{P}_{\mathrm{RL}}(l) = \frac{1}{\hat{\lambda}}\exp\left(-\frac{l}{\hat{\lambda}}\right)\left\{1 - \frac{1}{2(p_{0\mathrm{d}} + p_{5\mathrm{d}})}\sum_{j=0,5} p_{j\mathrm{d}}\left[\mathrm{erfc}(\sqrt{\alpha}\beta_j) + \sqrt{\frac{\alpha}{\pi}}\exp(-\alpha\beta_j^2)\right]\right\}. \tag{S33}$$

Furthermore, if $l^2 \gg \delta^2$ (this implies $\alpha\beta_j^2 \gg 1$) is satisfied, the quantity in the square brackets in this equation is much smaller than unity. Therefore, the run length distribution of the highly processive heterodimeric motor is well approximated by

$$\hat{P}_{\mathrm{RL}}(l) \approx \frac{1}{\hat{\lambda}} \exp\left(-\frac{l}{\hat{\lambda}}\right). \tag{S34}$$

Eqs. S31 and S34 correspond to Eqs. 8 and 7 in the main text, respectively.

#### S2-2 Run time of heterodimeric motor

To obtain the run time distribution of the heterodimeric motor, we introduce the following rate constants which approximate the transitions $1 \to 2 \to 3 \to 4 \to 5$ and transitions $6 \to 7 \to 8 \to 9 \to 0$, respectively:

$$\begin{aligned} L_1 &= \left(\frac{1}{l_1} + \frac{1}{l_2} + \frac{1}{l_3} + \frac{1}{l_4}\right)^{-1}, \\ L_6 &= \left(\frac{1}{l_6} + \frac{1}{l_7} + \frac{1}{l_8} + \frac{1}{l_9}\right)^{-1}. \end{aligned} \tag{S35}$$

We assume a four-state model governed by $l_0$, $l_{0\mathrm{d}}$, $L_1$, $l_5$, $l_{5\mathrm{d}}$, and $L_6$ shown in Fig. S1B, in which the motor starts running in state 0 at time $t = 0$. Let $Q_i$ be the probability that the motor is in state $i$ $(i = 0, 1, 5, 6)$ on the microtubule and we have the following master equations:

$$\begin{aligned} \frac{dQ_{0(t)}}{dt} &= L_6 Q_6(t) - (l_0 + l_{0\mathrm{d}}) Q_0(t), \\ \frac{dQ_{1(t)}}{dt} &= l_0 Q_0(t) - L_1 Q_1(t), \\ \frac{dQ_{5(t)}}{dt} &= L_1 Q_1(t) - (l_5 + l_{5\mathrm{d}}) Q_5(t), \\ \frac{dQ_{6(t)}}{dt} &= l_5 Q_5(t) - L_6 Q_6(t). \end{aligned} \tag{S36}$$

The solution to these coupled differential equations can be expressed as

$$Q_i(t) = \sum_{j=0}^{3} G_{ij} \exp(-\beta_j t) \qquad (i = 0, 1, 5, 6), \tag{S37}$$

where $\beta_j$ $(j = 0, 1, 2, 3)$ are the solutions to the quartic equation

$$(\beta - l_0 - l_{0\mathrm{d}})(\beta - L_1)(\beta - l_5 - l_{5\mathrm{d}})(\beta - L_6) = l_0 L_1 l_5 L_6 \tag{S38}$$

for $\beta$, and $G_{ij}$ are determined by

$$\begin{aligned} &(\beta_j - l_0 - l_{0\mathrm{d}}) G_{0j} + L_6 G_{6j} = 0, \\ &l_0 G_{0j} + (\beta_j - L_1) G_{1j} = 0, \\ &L_1 G_{1j} + (\beta_j - l_5 - l_{5\mathrm{d}}) G_{5j} = 0, \\ &l_5 G_{5j} + (\beta_j - L_6) G_{6j} = 0 \end{aligned} \qquad (j = 0, 1, 2, 3) \tag{S39}$$

together with the initial conditions $Q_0(0) = 1$ and $Q_1(0) = Q_5(0) = Q_6(0) = 0$.

The probability $Q(t)$ of the motor being on the microtubule at time $t$ is given as $Q = Q_0 + Q_1 + Q_5 + Q_6$, and from Eq. S37 we have

$$Q(t) = \sum_{j=0}^{3} S_j \exp(-\beta_j t), \tag{S40}$$

where

$$S_j = G_{0j} + G_{1j} + G_{5j} + G_{6j}. \tag{S41}$$

The run time distribution $\hat{P}_{\mathrm{RT}}(t)$ of the heterodimeric motor is obtained as follows:

$$\hat{P}_{\mathrm{RT}}(t) = -\frac{d}{dt}Q(t) = \sum_{j=0}^{3} S_j \beta_j \exp(-\beta_j t). \tag{S42}$$

If the heterodimer is highly processive, we can simplify Eq. S42 as follows. For the highly processive heterodimeric motor, we have small parameters

$$\epsilon_0 \equiv l_{0\mathrm{d}}/l_0 \quad \text{and} \quad \epsilon_5 \equiv l_{5\mathrm{d}}/l_5. \tag{S43}$$

These parameters will collectively be denoted by $\epsilon$. We rewrite Eq. S38 as

$$\beta^4 - A\beta^3 + B\beta^2 - C\beta + D = 0, \tag{S44}$$

where positive constants $A$, $B$, $C$, and $D$ can be expressed in terms of the rate constants. It is noted that

$$D = (\epsilon_0 + \epsilon_5 + \epsilon_0\epsilon_5) l_0 L_1 l_5 L_6 \tag{S45}$$

is of the first order in the small parameters $\epsilon$, which may be expressed as $D = O(\epsilon)$. On the other hand, $A$, $B$ and $C$ are of the zero-th order in $\epsilon$. From this fact it can be deduced that one of the four solutions $\beta_j$ ($j = 0, 1, 2, 3$) to Eq. S44, which will be labeled as $\beta_0$, is of the first order and the others, $\beta_j$ ($j \neq 0$), are of the zero-th order in $\epsilon$:

$$\beta_0 = O(\epsilon), \quad \beta_j = O(\epsilon^0) \quad (j \neq 0). \tag{S46}$$

Then, the first three terms in Eq. S44 are of the second or higher order in the case of $\beta = \beta_0$, and the approximate result $\beta_0 \approx D/C$ is obtained; keeping only the leading order terms, we have

$$\beta_0 \approx \frac{\tau_0 l_{0\mathrm{d}} + \tau_5 l_{5\mathrm{d}}}{\sum_{i=0}^{9} \tau_i}, \tag{S47}$$

where $\tau_i = 1/l_i$ ($0 \leqq i \leqq 9$), which means the lifetime of state $i$. The coefficients $S_j$ in Eq. S40 can be approximately obtained as follows. Summing up the four equations in Eq. S39, we obtain

$$\beta_j \left(G_{0j} + G_{1j} + G_{5j} + G_{6j}\right) = l_{0\mathrm{d}} G_{0\mathrm{d}} + l_{5\mathrm{d}} G_{5\mathrm{d}}. \tag{S48}$$

Note that the right-hand side of this equation is the first order in $\epsilon$ and remember that $\beta_j = O(\epsilon^0)$ for $j \neq 0$. Therefore, Eq. S48 togethr with Eq. S41 implies

$$S_j = O(\epsilon) \quad (j \neq 0). \tag{S49}$$

From this result and the initial condition $Q(0) = 1$, which reads $\sum_{j=0}^{3} S_j = 1$ according to Eq. S40, we obtain

$$S_0 = 1 + O(\epsilon). \tag{S50}$$

Now, from Eqs. S46, S49, and S50 we see that $S_j\beta_j$ in Eq. S42 are of $O(\epsilon)$ independent of $j$. However, $\exp(-\beta_j t)$ for $j \neq 0$ decays with increasing $t$ much faster than that for $j = 0$ due to Eq. S46. Therefore, the run time distribution $\hat{P}_{\mathrm{RT}}(t)$ for somewhat large $t$ and the mean run time $\tau_{\mathrm{run}}$ of the highly processive heterodimeric motor are approximately expressed as:

$$\hat{P}_{\mathrm{RT}}(t) = \beta_0 \exp(-\beta_0 t), \tag{S51}$$

$$\tau_{\mathrm{run}} = \frac{1}{\beta_0}, \tag{S52}$$

which correspond to Eqs. (9) and (10) in the main text, respectively.

#### S2-3 Inspection of the model for heterodimeric motor

Our calculations for heterodimers are demonstrated in Fig. S3 using the wt/R254Q heterodimer as an example. If the detaching rates of the heads are different, as in the case of the wt/R254Q heterodimer, and one of the heads binds to the microtubule first with higher probability than the other, then the run length distribution will show undulating behavior with large and small peaks alternating at even- and odd-integer multiples of the step size (Figs. S3A and B). However, if the spatial resolution is significantly larger than the step size and the motor exhibits high processivity, such peaks become almost invisible, and the distribution can be approximated by a single exponential, except in a narrow region near the zero of the run length (Fig. S3C).

The run time distribution is also affected by which head binds to the microtubule first. Supposing that head $X$ ($X$ = A or B) is the first one, we have $\hat{P}_{\mathrm{RT}}(0) = k^{X}_{0\mathrm{d}}$. However, for a highly processive motor, the motor quickly reaches a steady state in the stepping cycle, and the distribution approaches the exponential function in Eq. S51, which is independent of the initial condition (Fig. S3D). At the time resolution of 100 ms in this study, it is impossible to observe this transient behavior. Therefore, the distribution can be approximately described by a single exponential function (Fig. S3D).

### S3 Coordinated head model

In a saturated ADP solution, the motor consistently exists in a vulnerable one-head-bound state (state 0). If the detachment of the motor from this state is a single-rate-limiting process with a rate of $k_{0\mathrm{d}}$, the dwell time distribution for this state can be described as:

$$P_{\mathrm{DT}}(t) = k_{0\mathrm{d}} \exp(-k_{0\mathrm{d}} t). \tag{S53}$$

Now, we decompose state 0 into two states, $0'$ and $0''$. We assume that the motor can only detach from state $0'$, and that the transitions between these states occur reversibly with rates $k_{0'}$ and $k_{0''}$ as shown in Fig. S1C. In state $0'$, there are no interactions between the tethered head and the microtubule, while in state $0''$, there are interactions between them (Fig. 5A in the main text). Since the diagram in Fig. S1C is similar to the one in Fig. S1A for the homodimeric motor in the saturated ATP solution, the dwell time distribution for this modified model can be derived in a similar way to the run time of the homodimeric motor in S1-2. It is expected that $k_{0'}$ and $k_{0''}$ are much faster than $k_{0'\mathrm{d}}$. Therefore, we can adopt the approximate expressions, Eq. S21–S23, in the case of the highly processive motor. In this way, we obtain the modified dwell time distribution in high ADP concentration as follows:

$$P_{\mathrm{DT}}(t) \approx \frac{k_{0''} k_{0'\mathrm{d}}}{k_{0'} + k_{0''}} \exp\left(-\frac{k_{0''} k_{0'\mathrm{d}}}{k_{0'} + k_{0''}} t\right). \tag{S54}$$

Comparing Eq. S53 with Eq. S54, we obtain

$$k_{0\mathrm{d}} \approx r k_{0'\mathrm{d}}, \tag{S55}$$

where,

$$r = \frac{k_{0''}}{k_{0'} + k_{0''}}. \tag{S56}$$

If $k_{0'}$ and $k_{0''}$ are much faster than the other constant rates in the stepping cycle, the expression for $k_{0\mathrm{d}}$ in Eq. S55 allows us to use the kinetic diagram for the homodimer model in Fig. S1A and heterodimer model in Fig. S1B as they are, even in the presence of the interaction between the tethered head and the microtubule of the type considered here.

### S4 Simplified expression

It is explained in RESULTS of the main text that for the independent head model of heterodimeric motors, the mean run length $\hat{\lambda}$ and run time $\hat{\tau}_{\mathrm{run}}$ can be described by the rate constants of the parent homodimeric motors, as seen in Eqs. (16)–(19) of the main text. Here, we show that $\hat{\lambda}$ and $\hat{\tau}_{\mathrm{run}}$ can be simplified if the tethered head attachment is ruled by the microtubule-bound head (referred to as model I). For this model, we have the mean run length and run time of the heterodimeric motor, which correspond to Eqs. 16 and 18 of the main text, respectively, as follows:

$$\hat{\lambda} = \frac{2a k_0^{\mathrm{A}} k_0^{\mathrm{B}}}{k_0^{\mathrm{A}} k_{0\mathrm{d}}^{\mathrm{B}} + k_0^{\mathrm{B}} k_{0\mathrm{d}}^{\mathrm{A}}}, \tag{S57}$$

$$\hat{\tau}_{\mathrm{run}} = \frac{(\tau_{\mathrm{step}}^{\mathrm{A}} + \tau_{\mathrm{step}}^{\mathrm{B}}) k_0^{\mathrm{A}} k_0^{\mathrm{B}}}{k_0^{\mathrm{A}} k_{0\mathrm{d}}^{\mathrm{B}} + k_0^{\mathrm{B}} k_{0\mathrm{d}}^{\mathrm{A}}}, \tag{S58}$$

where the superscripts A and B indicate the types of homodimeric motors constituting the heterodimeric motor under consideration. Noting that the mean run length and run time of the homodimeric motor $X$ ($X$ = A or B) are given by Eqs. S10 and S22, respectively, i.e.,

$$\lambda^{X} = \frac{a k_0^{X}}{k_{0\mathrm{d}}^{X}}, \tag{S59}$$

$$\tau_{\mathrm{run}}^{X} = \frac{\tau_{\mathrm{step}}^{X} k_0^{X}}{k_{0\mathrm{d}}^{X}}, \tag{S60}$$

we can rewrite Eqs. S57 and S58 as

$$\hat{\lambda} = \frac{2\lambda^{A}\lambda^{B}}{\lambda^{A}+\lambda^{B}}, \tag{S61}$$

$$\hat{\tau}_{\text{run}} = \frac{\lambda^{A}\tau_{\text{run}}^{B}+\lambda^{B}\tau_{\text{run}}^{A}}{\lambda^{A}+\lambda^{B}}. \tag{S62}$$

The expressions in Eq. S61 and S62 look much simpler than Eqs. S57 and S58. Importantly, they demonstrate that we do not need all the details of the rate constants of the homodimeric motors to infer the run length and run time of the heterodimeric motor: only the run lengths and run times of the homodimeric motors are necessary. We also note that for model II, in which the tethered attachment is ruled by its own head, the run length and run time of the heterodimeric motor cannot be simply expressed in terms of those of the homodimeric motors.

### S5 Purification of recombinant KIF1A

BL21(DE3) cells transformed with KIF1A(1–393)::LZ::mScarlet-I::Strep plasmid were cultured in LB supplemented with kanamycin at 37 °C. Competent cells were prepared using a Mix & Go kit (Zymogen). The competent cells were further transformed with KIF1A(1-393)::LZ::His plasmid and selected on LB agar supplemented with ampicillin and kanamycin. Colonies were picked and cultured in 10 ml LB medium supplemented with ampicillin and kanamycin overnight. Next morning, 5 ml of the medium was transferred to 1000 ml 2.5 YT (20 g/L Tryptone, 12.5 g/L Yeast Extract, 6.5 g/L NaCl) supplemented with 10 mM phosphate buffer (pH 7.4), ampicillin and kanamycin in a 2 L flask and shaken at 37 °C. Two flasks were routinely prepared. When OD600 reached 0.6, flasks were cooled in ice-cold water for 30 min. Then, 23.8 mg IPTG was added to each flask. Final concentration of IPTG was 0.2 mM. Flasks were shaken at 18 °C overnight. Next day, bacteria expressing recombinant proteins were pelleted by centrifugation (3000 g, 10 min, 4 °C), resuspended in PBS and centrifuged again (3000 g, 10 min, 4 °C). Pellets were resuspended in protein buffer (50 mM Hepes, pH 7.4, 150 mM $KCH_3COO$, 2 mM $MgSO_4$, 1 mM EGTA, 10 % glycerol) supplemented with Phenylmethylsulfonyl fluoride (PMSF). Bacteria were lysed using a French Press G-M (Glen Mills, NJ, USA) as described by the manufacturer. Lysate was obtained by centrifugation (75,000 g, 20 min, 4 °C). Lysate was loaded on Streptactin-XT resin (IBA Lifesciences, Göttingen) (bead volume: 2 ml). The resin was washed with 40 ml wash buffer (50 mM Hepes, pH 8.0, 450 mM $KCH_3COO$, 2 mM $MgSO_4$, 1 mM EGTA, 10 % glycerol). Protein was eluted with 40 ml protein buffer supplemented with 50 mM biotin. Eluted solution was then loaded on TALON resin (Takara Bio Inc., Kusatsu, Japan)(bead volume: 2 ml). The resin was washed with 40 ml wash buffer and eluted with 40 ml protein buffer supplemented with 500 mM imidazole. Eluted solution was concentrated using an Amicon Ultra 15 (Merck) and then separated on an NGC chromatography system (Bio-Rad) equipped with a Superdex 200 Increase 10/300 GL column (Cytiva). The heterodimers were recovered from the same peak fractions (Fig. S4C), where the ratio between two subunits calculated from band intensities and molecular weight was about 1:1 (Fig. S4D). After concentration using an Amicon Ultra 4 (Merck), they were aliquoted and snap frozen in liquid nitrogen.

### S6 TIRF single-molecule motility assays

TIRF assays were performed as described (4, 5). Tubulin was purified from porcine brain as described (6). Tubulin was labeled with Biotin-PEG2-NHS ester (Tokyo Chemical Industry, Tokyo, Japan) and AZDye647 NHS ester (Fluoroprobes, Scottsdale, AZ, USA) as described (7). To polymerize Taxol-stabilized microtubules labeled with biotin and AZDye647, 30 $\mu$M unlabeled tubulin, 1.5 $\mu$M biotin-labeled tubulin and 1.5 $\mu$M AZDye647-labeled tubulin were mixed in BRB80 buffer supplemented with 1 mM GTP and incubated for 40 min at 37°C. Then, an equal amount of BRB80 supplemented with 40 $\mu$M taxol was added and further incubated for more than 40 min. The solution was loaded on BRB80 supplemented with 300 mM sucrose and 20 $\mu$M taxol and ultracentrifuged at 100,000 g for 10 min at 30°C. The pellet was resuspended in BRB80 supplemented with 20 $\mu$M taxol. Glass chambers were prepared by acid washing as previously described (8). Glass chambers were coated with PLL-PEG-biotin (SuSoS, Dübendorf, Switzerland). Polymerized microtubules were flowed into streptavidin adsorbed flow chambers and allowed to adhere for 5–10 min. Unbound microtubules were washed away using assay buffer [90 mM Hepes pH 7.4, 100 mM $KCH_3COO$, 2 mM $Mg(CH_3COO)_2$, 1 mM EGTA, 10% glycerol, 0.1 mg/ml biotin–BSA, 0.2 mg/ml kappa- casein, 0.5% Pluronic F127, 2 mM ATP, and an oxygen scavenging system composed of PCA/PCD/Trolox. Purified motor protein was diluted to indicated concentrations in the assay buffer. Then, the solution was flowed into the glass chamber. All assays were performed at 25 °C . An ECLIPSE Ti2-E microscope equipped with a CFI Apochromat TIRF 100XC Oil objective lens, an Andor iXion life 897 camera and a Ti2-LAPP illumination system (Nikon, Tokyo, Japan) was used to observe single molecule motility. NIS-Elements AR software ver. 5.2 (Nikon) was used to control the system. For quantification, kymographs were made by ImageJ software. We analyzed the motor that bound to microtubules within the observation time window. Unidirectional lines greater than 3 pixels along x axis (corresponding to approximately 480 nm) were counted as processive runs and the other signals were not counted.

### S7 Graph preparation

Graphs were prepared using Python and Graph Pad Prism version 9. They were exported in the PNG format, and aligned by Adobe Illustrator 2021.


### Supporting Table

Table S1: Plasmid list

| Plasmid | Description | Backbone |
|---|---|---|
| pSN643 | KIF1A(1-393)::LZ::mScarlet-I::Strep tag | pET28a |
| pSN655 | KIF1A(1-393)(V8M)::LZ::mScarlet-I::Strep tag | pET28a |
| pSN657 | KIF1A(1-393)(R254Q)::LZ::mScarlet-I::Strep tag | pET28a |
| pSN826 | KIF1A(1-393)(T258M)::LZ::mScarlet-I::Strep tag | pET28a |
| pSN787 | KIF1A(1-393)(R350G)::LZ::mScarlet-I::Strep tag | pET28a |
| pSN672 | KIF1A(1-393)::LZ::His tag | pET21a |
| pSN744 | KIF1A(1-393)(V8M)::LZ::His tag | pET21a |
| pSN686 | KIF1A(1-393)(R254Q)::LZ::His tag | pET21a |
| pTK2 | KIF1A(1-393)(T258M)::LZ::His tag | pET21a |
| pTK3 | KIF1A(1-393)(R350G)::LZ::His tag | pET21a |

### Supporting Figures

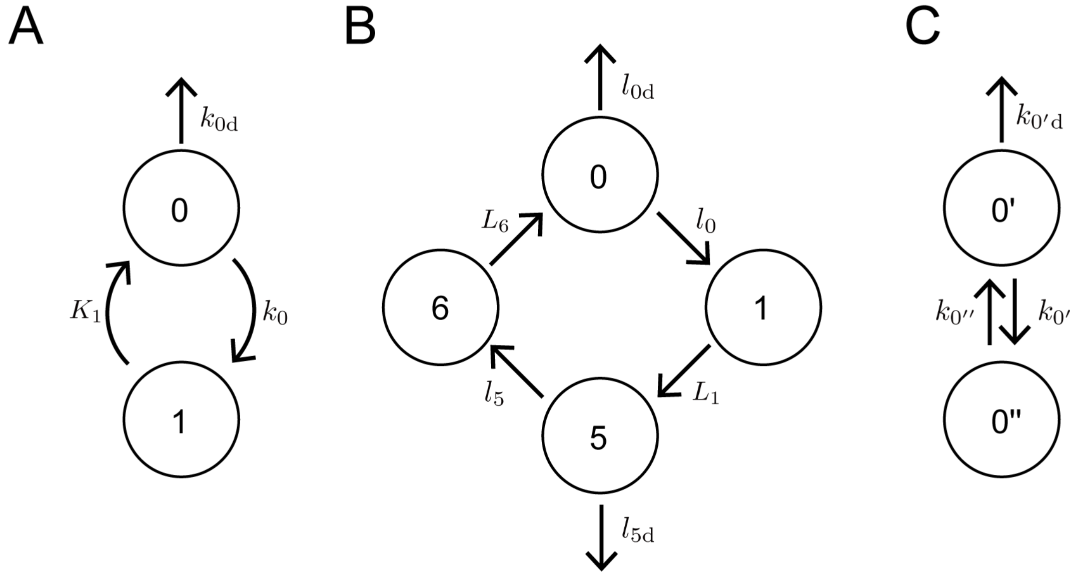


Figure S1: Kinetic diagram for homodimer and heterodimer models. Symbols $k$, $K$, $l$, and $L$ indicate rate constants. (A) Two state-model for homodimers, (B) Four-state model for heterodimers, and (C) Two state-model for homodimers in ADP solution.

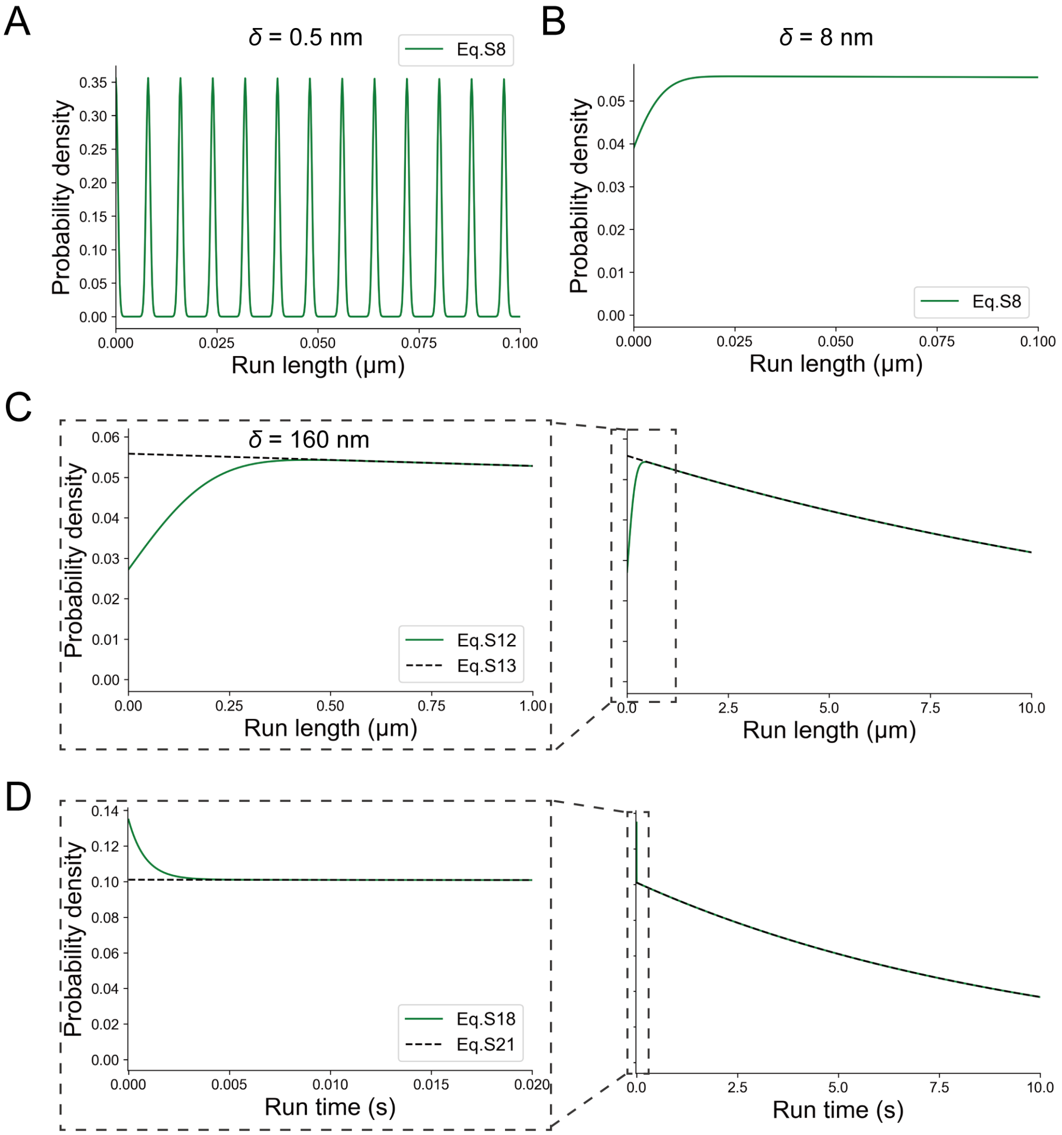


Figure S2: Theoretical predictions are shown for (A-C) run length distributions observed at various spatial resolutions and (D) run time distribution of KIF1A homodimer. The parameters $k_0^{\mathrm{wt}} = 302\ \mathrm{s}^{-1}$, $k_{0d}^{\mathrm{wt}} = 0.13\ \mathrm{s}^{-1}$, $\tau_{\mathrm{step}}^{\mathrm{wt}} = 4.4 \times 10^{-3}$ s, and $K_1^{\mathrm{wt}} = (\tau_{\mathrm{step}}^{\mathrm{wt}} + 1/k_0^{\mathrm{wt}})^{-1} = 904\ \mathrm{s}^{-1}$ corresponding to wt/wt homodimer are used. The solid lines in (A) and (B) are plotted by using Eq. S8 with the aforementioned values, and spatial resolutions of $\delta = 0.5$ nm and $\delta = 8$ nm, respectively. The solid and dashed lines in (C) are plotted by using Eqs. S12 and S13, respectively, with the aforementioned values and $\delta = 160$ nm. The solid and dashed lines in (D) are plotted by using Eqs. S18 and S21, respectively, with the aforementioned values. The graphs on the right in (C) and (D) show a larger area than the graphs on the left.

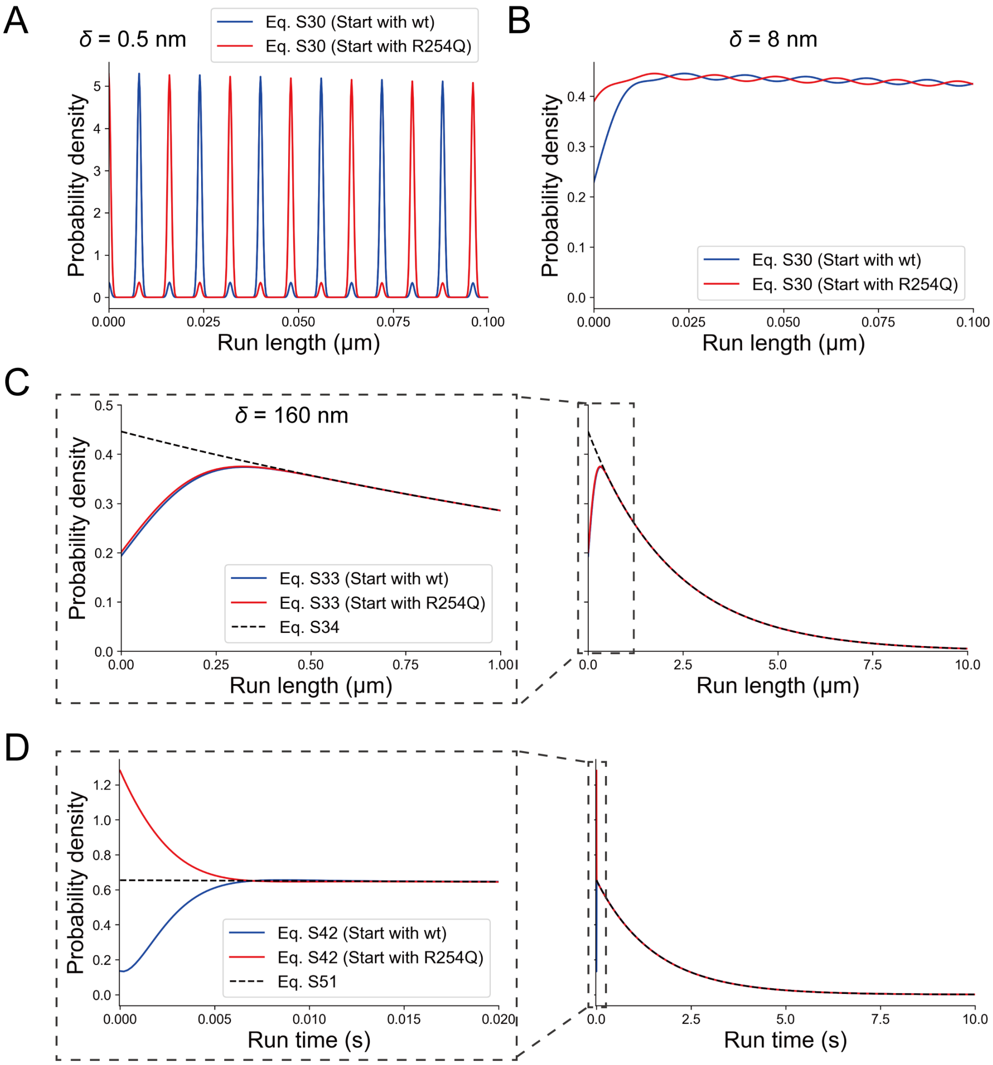


Figure S3: Theoretical predictions are shown for (A-C) run length distributions observed at various spatial resolutions and (D) run time distribution of KIF1A heterodimer. The run length and run time distributions of wt/R254Q heterodimers are described by the independent head model, in which the tethered head attachment is ruled by the microtubule-bound head (referred to as model I in the main text) using the following homodimer parameters: $k_0^{\mathrm{wt}} = 302\ \mathrm{s}^{-1}$, $k_{0\mathrm{d}}^{\mathrm{wt}} = 0.13\ \mathrm{s}^{-1}$, $\tau_{\mathrm{step}}^{\mathrm{wt}} = 4.4 \times 10^{-3}$ s, and $K_1^{\mathrm{wt}} = (\tau_{\mathrm{step}}^{\mathrm{wt}} + 1/k_0^{\mathrm{wt}})^{-1} = 904\ \mathrm{s}^{-1}$ for wt/wt homodimer, and $k_0^{\mathrm{R254Q}} = 192\ \mathrm{s}^{-1}$, $k_{0\mathrm{d}}^{\mathrm{R254Q}} = 1.28\ \mathrm{s}^{-1}$, $\tau_{\mathrm{step}}^{\mathrm{R254Q}} = 6.5 \times 10^{-3}$ s, and $K_1^{\mathrm{R254Q}} = (\tau_{\mathrm{step}}^{\mathrm{R254Q}} + 1/k_0^{\mathrm{R254Q}})^{-1} = 790\ \mathrm{s}^{-1}$ for R254Q/R254Q homodimer. We assume that $K_1$, $L_1$, and $L_6$ are approximately determined by the ADP release rate from the front head in a two-head-bound state, which is the slowest reaction among all the reactions included in $K_1$, $L_1$, and $L_6$, respectively (2). The solid blue line represents the result under the condition in which the heterodimer initiates the run with the wild-type head bound to the microtubule. The parameters are set as $l_0 = k_0^{\mathrm{wt}}$, $l_{0\mathrm{d}} = k_{0\mathrm{d}}^{\mathrm{wt}}$, $L_1 \approx K_1^{\mathrm{wt}}$, $l_5 = k_0^{\mathrm{R254Q}}$, $l_{5\mathrm{d}} = k_{0\mathrm{d}}^{\mathrm{R254Q}}$, and $L_6 \approx K_1^{\mathrm{R254Q}}$. The red solid line represents the result under the condition in which the heterodimer initiates the run with the R254Q head bound to the microtubule. The parameters are set as $l_0 = k_0^{\mathrm{R254Q}}$, $l_{0\mathrm{d}} = k_{0\mathrm{d}}^{\mathrm{R254Q}}$, $L_1 \approx K_1^{\mathrm{R254Q}}$, $l_5 = k_0^{\mathrm{wt}}$, $l_{5\mathrm{d}} = k_{0\mathrm{d}}^{\mathrm{wt}}$, and $L_6 \approx K_1^{\mathrm{wt}}$. The black dashed line represents the approximate solution, which is not affected by which head the motor starts the run with. The parameters are set as $l_0 = k_0^{\mathrm{wt}}$, $l_{0\mathrm{d}} = k_{0\mathrm{d}}^{\mathrm{wt}}$, $l_5 = k_0^{\mathrm{R254Q}}$, $l_{5\mathrm{d}} = k_{0\mathrm{d}}^{\mathrm{R254Q}}$, and $\tau_{2\mathrm{step}} = \tau_{\mathrm{step}}^{\mathrm{wt}} + \tau_{\mathrm{step}}^{\mathrm{R254Q}}$. The solid lines in (A) and (B) are plotted by using Eq. S30 with the aforementioned values, and spatial resolutions of $\delta = 0.5$ nm and $\delta = 8$ nm, respectively. The solid and dashed lines in (C) are plotted by using Eqs. S33 and S34, respectively, with the aforementioned values and $\delta = 160$ nm. The solid and dashed lines in (D) are plotted by using Eqs. S42 and S51, respectively, with the aforementioned values. The graphs on the right in (C) and (D) show a larger area than the graphs on the left.

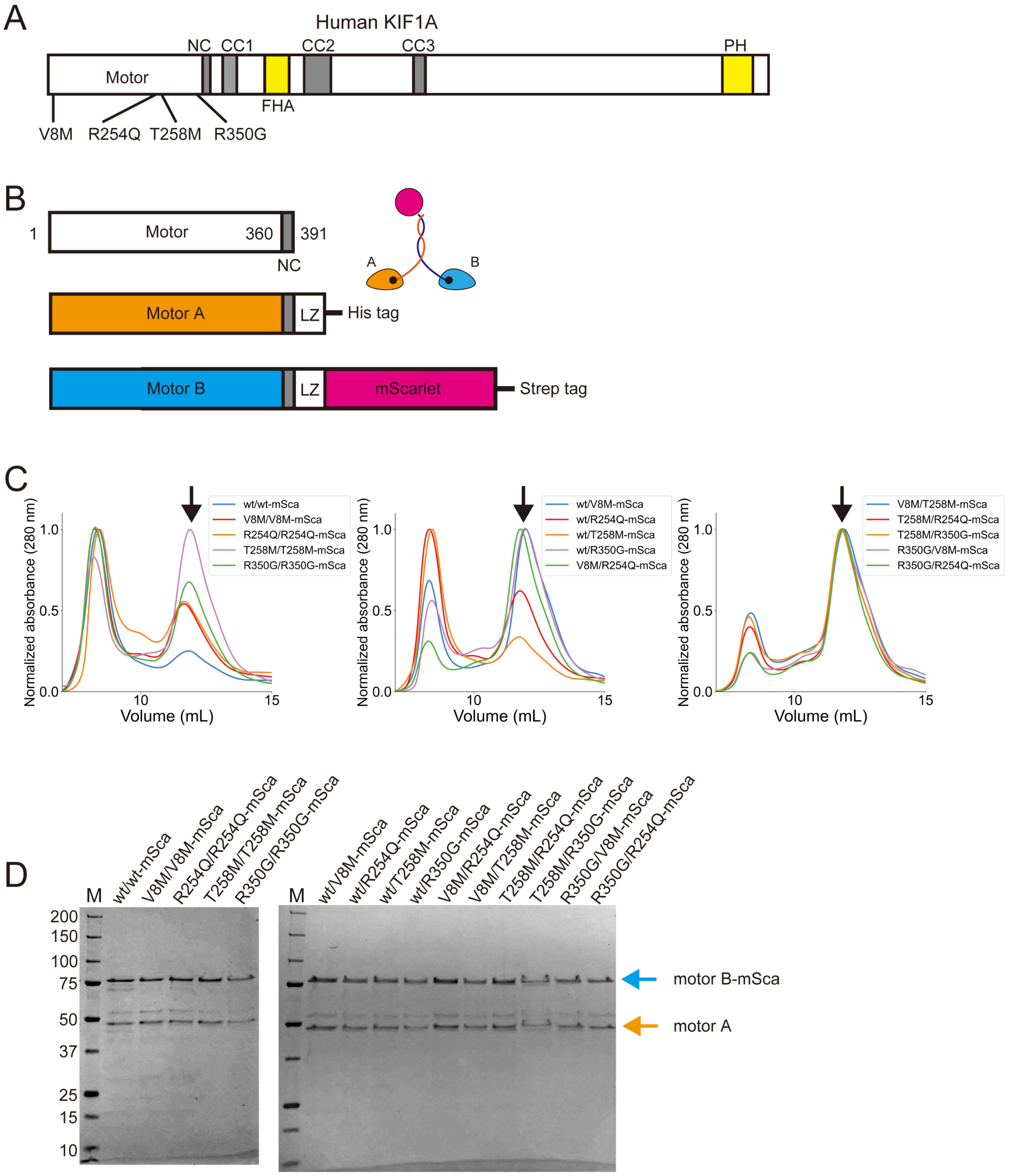


Figure S4: Purification of recombinant KIF1A (A) Schematic drawing of the domain organization of KIF1A motor protein. NC, neck coiled-coil domain. CC1, Coiled-coil 1 domain. FHA, Forkhead-associated domain. CC2, Coiled-coil 2 domain. CC3, Coiled-coil 3 domain. The four KAND mutations analyzed in this study are indicated. (B) Schematic drawing of the recombinant KIF1A dimer analyzed. The KIF1A dimer is composed of KIF1A(1-393)::His::LZ, referred to as motor A, and KIF1A(1-393)::Strep::LZ::mScarlet, referred to as motor B-mSca. (C) Results of gel filtration. The recombinant KIF1A was recovered from the peak fraction shown by arrow heads. (D) The purified KIF1A dimers composed of motor A and motor B-mSca were separated by SDS-PAGE and detected by Coomassie brilliant blue staining. M represents marker. Numbers on the left indicate the molecular weight (kDa). Orange and blue arrows indicate motor A and motor B-mSca, respectively.

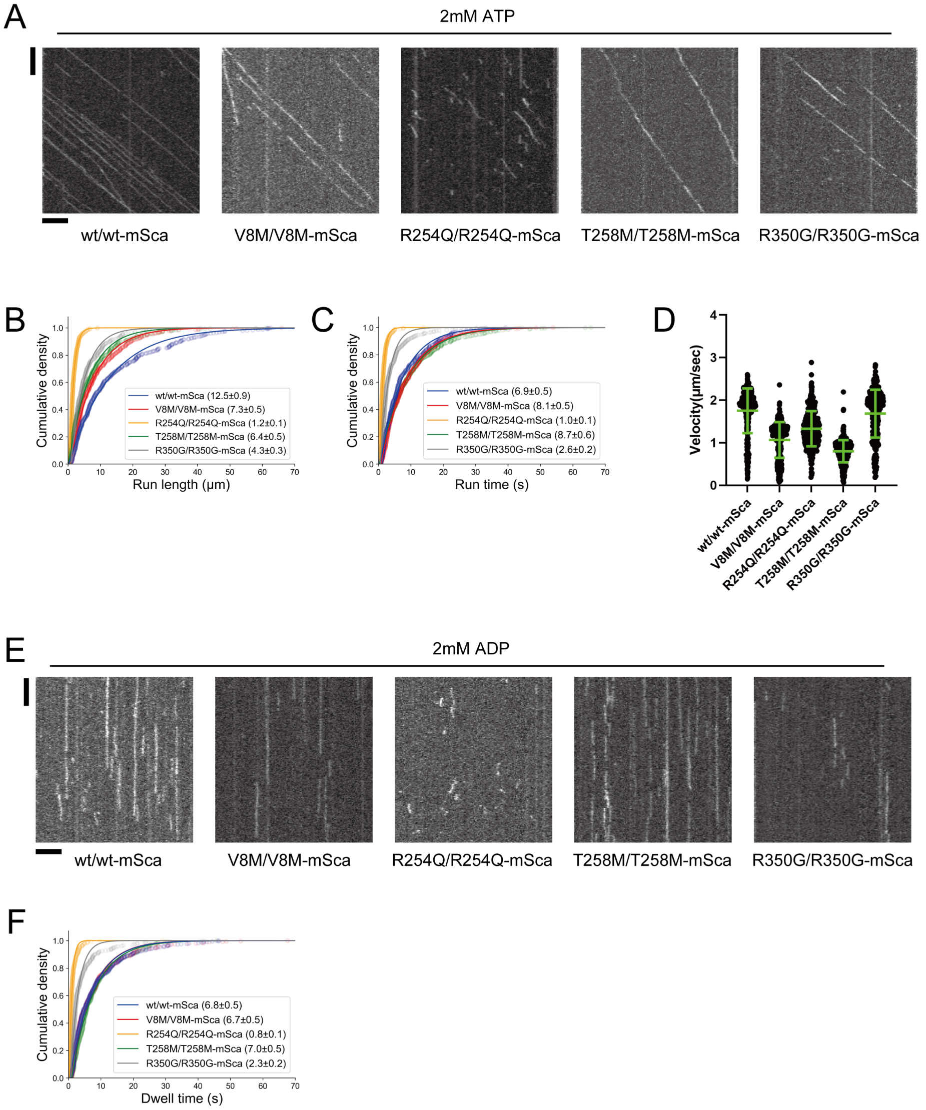


Figure S5: The single-molecule behavior of KIF1A homodimers. (A) Representative kymographs showing the motility of each homodimer in the presence of 2mM ATP. Vertical and horizontal bars represent 5 s and 5 $\mu$m, respectively. (B) Cumulative density plots showing the run lengths of each homodimer in the presence of 2mM ATP. n = 377 (wt/wt-mSca), 418 (V8M/V8M-mSca), 385 (R254Q/R254Q-mSca), 405 (T258M/T258M-mSca), 335 (R350G/R350G-mSca) molecules. Values are fit by Eq. 12 in the main text with bootstrapping error. (C) Cumulative density plots showing the run times of each homodimer in the presence of 2mM ATP. n = 377 (wt/wt-mSca), 418 (V8M/V8M-mSca), 385 (R254Q/R254Q-mSca), 405 (T258M/T258M-mSca), 335 (R350G/R350G-mSca) molecules. Values are fit by Eq. 12 in the main text with bootstrapping error. (D) Dot plots showing the velocity of each homodimer in the presence of 2mM ATP. Each dot shows a single datum point. Green bars represent mean ± SD. n = 377 (wt/wt-mSca), 418 (V8M/V8M-mSca), 385 (R254Q/R254Q-mSca), 405 (T258M/T258M-mSca), 335 (R350G/R350G-mSca) molecules. (E) Representative kymographs showing the binding of each homodimer with microtubules in the presence of 2mM ADP. Vertical and horizontal bars represent 5 s and 5 $\mu$m, respectively. (F) Cumulative density plots showing the dwell times of each homodimer in the presence of 2mM ADP. n = 354 (wt/wt-mSca), 357 (V8M/V8M-mSca), 318 (R254Q/R254Q-mSca), 290 (T258M/T258M-mSca), 287 (R350G/R350G-mSca) molecules. Values are fit by Eq. 12 in the main text with bootstrapping error.

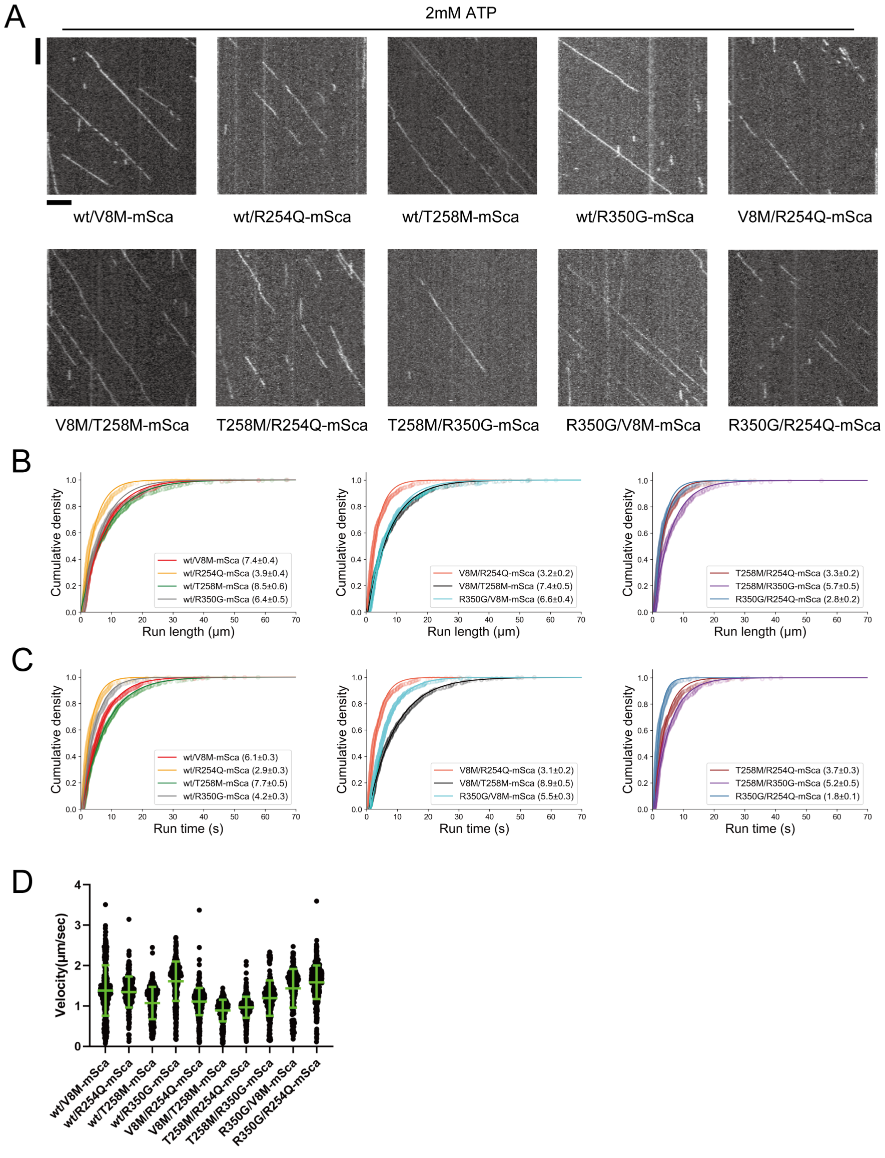


Figure S6: The single-molecule behavior of KIF1A heterodimers. (A) Representative kymographs showing the motility of each heterodimer in the presence of 2mM ATP. Vertical and horizontal bars represent 5 s and 5 $\mu$m, respectively. (B) Cumulative density plots showing the run times of each heterodimer in the presence of 2mM ATP. n = 500 (wt/V8M-mSca), 311 (wt/R254Q-mSca), 420 (wt/T258M-mSca), 398 (wt/R350G-mSca), 537 (V8M/R254Q-mSca), 330 (V8M/T258M-mSca), 480 (T258M/R254Q-mSca), 246 (T258M/R350G-mSca), 254 (R350G/V8M-mSca), 501 (R350G/R254Q-mSca) molecules. Values are fit by Eq. 12 in the main text with bootstrapping error. (C) Cumulative density plots showing the run lengths of each heterodimer in the presence of 2mM ATP. n = 500 (wt/V8M-mSca), 311 (wt/R254Q-mSca), 420 (wt/T258M-mSca), 398 (wt/R350G-mSca), 537 (V8M/R254Q-mSca), 330 (V8M/T258M-mSca), 480 (T258M/R254Q-mSca), 246 (T258M/R350G-mSca), 254 (R350G/V8M-mSca), 501 (R350G/R254Q-mSca) molecules. Values are fit by Eq. 12 in the main text with bootstrapping error. (D) Dot plots showing the velocity of each heterodimer in the presence of 2mM ATP. Each dot shows a single datum point. Green bars represent mean ± SD. n = 500 (wt/V8M-mSca), 311 (wt/R254Q-mSca), 420 (wt/T258M-mSca), 398 (wt/R350G-mSca), 537 (V8M/R254Q-mSca), 330 (V8M/T258M-mSca), 480 (T258M/R254Q-mSca), 246 (T258M/R350G-mSca), 254 (R350G/V8M-mSca), 501 (R350G/R254Q-mSca) molecules.

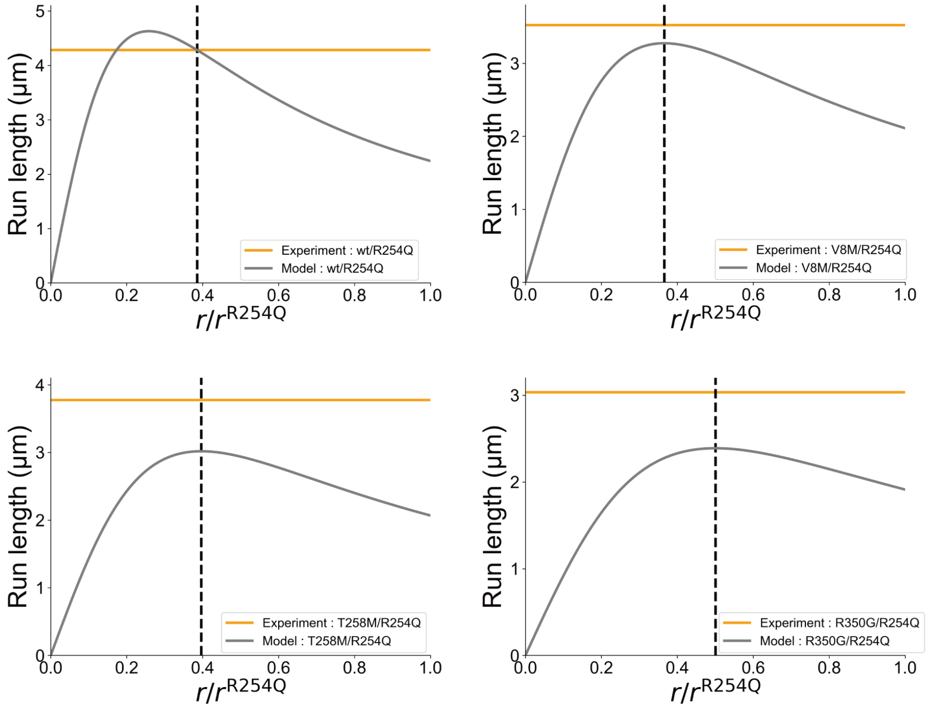


Figure S7: Determination of $r^{\mathrm{A}}/r^{\mathrm{R254Q}}$ for motor A with A ≠ R254Q. Horizontal orange solid line in each panel shows the experimental mean run length of heterodimer composed of motor A and R254Q. Gray solid line shows the dependence of the run length for dimer A/R254Q predicted by Eq. 24 in the main text. Black dashed lines show the determined values of $r^{\mathrm{A}}/r^{\mathrm{R254Q}}$ specific to each motor. If there are two intersections of orange and gray lines, the one satisfying $l_{0\mathrm{d}} < l_{5\mathrm{d}}$ was selected. If there are no intersections, the value that minimizes the distance between the two lines was selected. We obtained $r^{\mathrm{wt}}/r^{\mathrm{R254Q}} = 0.39$, $r^{\mathrm{V8M}}/r^{\mathrm{R254Q}} = 0.37$, $r^{\mathrm{T258M}}/r^{\mathrm{R254Q}} = 0.40$, and $r^{\mathrm{R350G}}/r^{\mathrm{R254Q}} = 0.50$.

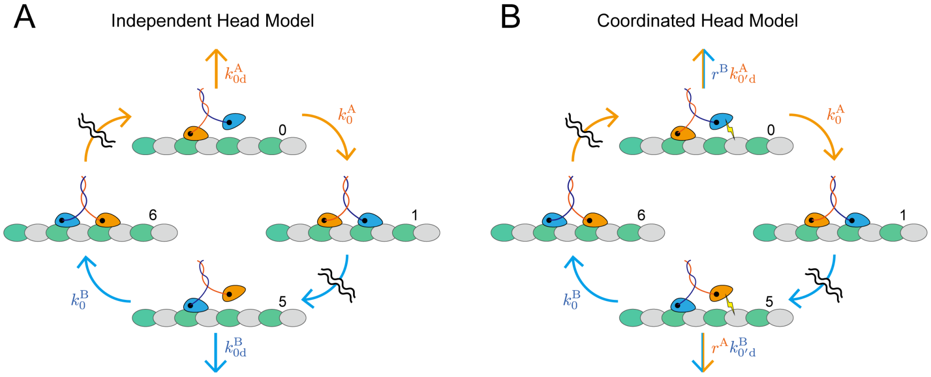


Figure S8: Comparison of (A) the independent head model and (B) the coordinated head model. States 2 – 4 and 7 – 9 in both models, not indicated in the diagrams, correspond to the states shown in Fig. 2 in the main text. Transitions from these states are governed by the head in which the corresponding reactions occur, as shown in Fig. 3A in the main text. The tethered head attachment (transitions 0 ⟶ 1 and 5 ⟶ 6) is determined by the microtubule-bound head, with a rate of $k_0^X$ ($X$ = A or B) in both models. In the independent head model, detachment from the microtubule is ruled solely by the microtubule-bound head, with a rate of $k_{0\mathrm{d}}^X$. However, in the coordinated head model, as the tethered head also contributes to the association between the motor and the microtubule, motor detachment is influenced by both the microtubule-bound head and the tethered head. $k_{0\mathrm{d}'}^X$ and $r^Y$ ($Y$ = A or B, and $Y \neq X$) are associated with the microtubule-bound head and the tethered head, respectively.
